# Reproducibility and reliability of Free-Water-corrected Diffusion Tensor Imaging of the brain: Revisited

**DOI:** 10.1101/2025.08.06.668995

**Authors:** Tomasz Pieciak, Guillem París, Santiago Aja-Fernández, Antonio Tristán-Vega

**Affiliations:** Laboratorio de Procesado de Imagen (LPI), ETSI Telecomunicación, Universidad de Valladolid, Valladolid, Spain

**Keywords:** diffusion tensor imaging, free-water, brain, white matter, reproducibility, reliability, separability

## Abstract

Diffusion tensor imaging (DTI) corrected for the free-water (FW) enables the separation of a hindered Gaussian-like profile from an isotropic component, which represents diffusion found in cerebrospinal and interstitial fluids within the extracellular space of grey and white matter. The assessment of the reproducibility and reliability properties of FW-corrected DTI is a crucial factor in demonstrating the potential clinical utility of this refinement, particularly considering the examinations across multiple medical centres. This paper explores the variability, reliability, and separability properties of free-water volume fraction (FWVF) and FW-corrected DTI-based measures in healthy human brain white matter using publicly available test-retest databases acquired in 1) intra-scanner, 2) intra-scanner longitudinal and 3) inter-scanner settings under varying acquisition schemes. Three different estimation techniques to retrieve the FW-corrected DTI parameters tailored to single- or multiple-shell diffusion-sensitizing magnetic resonance (MR) acquisitions are investigated: i) a direct optimization of bi-tensor signal representation in the variational framework, ii) the region contraction-based approach, and iii) the spherical means technique combined with a correction of diffusion-weighted MR signal prior to DTI estimation. We found the previous suggestion (Hum. Brain Mapp., 38(1), 2017, 10.1002/hbm.23350) that the FW correction to DTI in a single-shell diffusion-weighted MR acquisition improves the repeatability and reliability of DTI-based measures may be data- and methodology-dependent, and does not generalise to multiple-shell scenario. Our experiments have shown that the most reliable and repeatable/reproducible measures, while preserving a moderate separability property, are fractional anisotropy and axial diffusivity estimated in a multiple-shell variant under a combined FW-correction scheme. On the contrary, our results show evidence that the least reliable measures are the mean diffusivity estimated using any FW-correction procedure, as well as the FWVF parameter itself. These results can be used to establish the direction for selecting the most attractive FW-correction DTI scheme for clinical applications in terms of the variability-reliability-separability criterion.

## 1. Introduction

Diffusion-weighted magnetic resonance imaging (MRI) is a well-established medical modality that enables the non-invasive probing of random motion of water molecules *in vivo*, particularly in brain tissue (Le Bihan & Johansen-Berg, 2012). A common approach employed to represent the diffusion-weighted MR signal is single-compartment diffusion tensor imaging (DTI) by providing a set of unique quantitative measures summarizing the directional water diffusion process (Basser et al., 1994; Westin et al., 2002). Extending the standard DTI to the so-called bi-tensor representation has facilitated the separation of hindered diffusion depicted with a tensor-based Gaussian-like profile and the isotropic component, which illustrates the diffusion found in cerebrospinal and interstitial fluids within the extracellular space of grey and white matter (Pierpaoli & Jones, 2004; Pasternak et al., 2009). This separation is possible due to suitably selected numerical optimization schemes that enable to compute the directional DTI profile and the free-water volume fraction (FWVF), a scalar parameter illustrating the fitted isotropic fraction of the bi-tensor representation to the diffusion-sensitized MR signal. Such computations are possible both for single- (Pasternak et al., 2009) and multiple-shell acquisitions (Pasternak et al., 2012; Hoy et al., 2014; Bergmann et al., 2020; Tristán-Vega et al., 2022).

The FW-corrected DTI has been an essential tool in clinical applications, particularly the single-shell FW-corrected DTI, which is primarily employed in cognitive performance evaluation (Maillard et al., 2019), modelling neurodegenerative disorders such as the Parkinson’s (Ofori et al., 2015) or Alzheimer’s (Bergamino et al., 2021; Nakaya et al., 2022), brain ageing (Metzler-Baddeley et al., 2012; Chad et al., 2018), schizophrenia (Carreira Figueiredo et al., 2022) or detecting first-episode of psychosis (Lyall et al., 2018), as the evidence indicates the method improves the specificity of DTI measures (Bergamino et al., 2021; Chad et al., 2023) and reliability and accuracy of tractometry in brain ageing (Chang et al., 2024). However, the single-shell FW-corrected DTI by Pasternak et al. (2009) has been lately put into stake because it is unclear whether it accurately captures actual anatomy-related changes in the diffusivity profile of brain white matter. For instance, Golub et al. (2021) have shown that the single-shell FW-correction scheme can yield plausible results, but the optimization procedure heavily depends on the initialization strategy, potentially affecting method’s specificity. Correia et al. (2024) discovered a flattening effect of FW-corrected MD profiles with age and FW-corrected FA strong positive correlations with age in some regions, which seem not to be present in a multiple-shell scenario. The authors of these works advocate for considering the multiple-shell rather than single-shell tailored schemes once correcting the DTI for the FW component. As an example, the multiple-shell variant has explicitly demonstrated benefits over the single-shell in brain age predictions (Nemmi et al., 2022).

Although numerous FW-corrected DTI applications have already been demonstrated in a wide range of clinical scenarios, much less attention has been paid to the reproducibility and reliability of the measures obtained from this refinement. Until now, the intra-scanner repeatability (or more generally, the inter-scanner reproducibility) and reliability studies have concentrated primarily on the standard DTI-based metrics under specific acquisition conditions, such as variable magnetic strength fields (Grech-Sollars et al., 2015; Venkatraman et al., 2015; Jakab et al., 2017), multi-band acquisition scheme (Duan et al., 2015), and spatial resolution (Shahim et al., 2017; Zhong et al., 2024), considering particular cohorts like fetal brains (Jakab et al., 2017), neonates (Merisaari et al., 2019) or in older population (Laguna et al., 2020), in a longitudinal scenario (Boudreau et al., 2025), across multiple centres (Grech-Sollars et al., 2015) or even under scanner relocation (Melzer et al., 2020). Apart from providing detailed quantitative results, some other studies proposed vendor-agnostic sequences to reduce inter-scanner variabilities (Liu et al., 2024) or drawn conclusions on possible solutions to post-hoc improve the reproducibility or reliability of DTI-based parameters. For instance, Jakab et al. (2017) presented a negative impact of fetal head motion on DTI repeatability and suggested that this effect can be partially palliated with motion correction algorithms. Ades-Aron et al. (2024) have shown that proper denoising of diffusion-weighted MR data (either in complex or magnitude space) leads to a significant reduction in variability of DTI-based measures and increases statistical power for low signal-to-noise ratio voxels in intra- and inter-scanner, and inter-protocol studies. In the work by Albi et al. (2017), it has been suggested that suppressing the FW component in DTI using the single-shell approach by Pasternak et al. (2009) leads to reduced repeatability errors of standard metrics such as fractional anisotropy (FA) or mean diffusivity (MD) *on average* approximately at 1%pt^1^. However, taking a closer look at the experimental methodology, the work by Albi et al. (2017) may lead to more inquiries than answers, as different numerical optimization schemes have been arranged to compare the standard DTI-based measures and the FW-corrected ones, i.e., the linear least squares procedure *versus* the non-linear optimization in a variational framework. Besides, the work utilises a questionable single-shell-based FW correction scheme and uses only a test-retest database acquired in an intra-scanner scenario, i.e., it evaluates only the repeatability, not reproducibility across the centres.

In this paper, we revisit the study by Albi et al. (2017) and explore the repeatability and reproducibility, reliability and separability of FW-corrected DTI under three different approaches used to correct the DTI for the FW component, namely 1) a single-shell variational scheme by Pasternak et al. (2009), 2) the multiple-shell region contraction-based approach by Hoy et al. (2014), and 3) a prior correction of diffusion-weighted MR signal for the FW component estimated using the spherical means technique by Tristán-Vega et al. (2022), preceded by a standard DTI estimation from FW-corrected signal. Our study encompasses three databases acquired in i) intra-scanner, ii) intra-scanner longitudinal and iii) inter-scanner scenarios, as well as under various experimental setups. The results show evidence that the improved repeatability of the FW-corrected DTI in a single-shell scenario observed by Albi et al. (2017) may be data- and methodology-dependent, and does not generalize to tailored multiple-shell FW correction schemes, like Hoy et al. (2014). On the contrary, we have shown that the FW-corrected DTI in a multiple-shell scenario, using a region contraction-based technique, leads to systematic declines in repeatability/reproducibility and reliability compared to the standard DTI, as the number of degrees of freedom in the optimisation procedure is larger. Our study shows evidence that the most reliable and repeatable (and reproducible) measures are FA, AD (axial diffusivity) and RD (radial diffusivity) estimated from the standard DTI and among the FW-corrected DTI-based measures the FA and AD estimated from a previously corrected diffusion-weighted MR signal under a multiple-shell variant. In contrast, the least reliable and separable measure is the MD obtained from any FW correction approach, as well as the FWVF parameter itself, no matter whether estimated separately *via* the spherical means technique or jointly with the DTI.

## 2. Materials and methods

### 2.1. In silico data

We generate *in silico* data by composing the signals originating from cellular and free-water compartments, following the formulation:

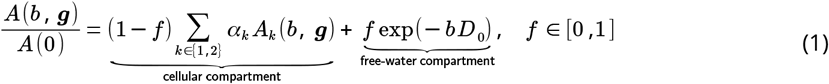

with ***D* = 3.0 × 10**^**−3**^ **mm**^**2**^**/ s** being the apparent diffusion coefficient for water at a temperature of approximately **37**^***°***^ **C**, *f* is the FWVF parameter, and ***A_k_* (*b***, ***g*)** is the cellular compartment integrating intra- and extra-axonal parts spherically convolved with the fibre orientation density function **Φ(*g*)**

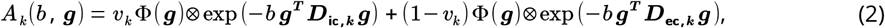

where the tensor ***D***_**ic**,***k***_ represents the intra-cellular part and is characterized with two perpendicular null eigenvalues,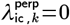, and the tensor ***D*** _**ec**,***k***_ is axis-symmetric with a non-zero perpendicular diffusivity 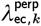. The cellular compartment defined in Eq. (1) can be represented using a single- or two-fibre bundle with partial fractions α_1_**+** α_2_**=1**. The signal given by Eq. (1) is then contaminated with a Rician noise (Aja-Fernández and Vegas-Sánchez-Ferrero, 2016; Pieciak et al., 2017), as follows ***S* (*b***, ***g*) =**|***A*(*b***, ***g*) + *N*** _**re**_ **+ *j*·*N*** _im_| with ***N*** _**re**_, ***N*_im_ ∼*N* (0**, *σ*^2^). For more information on synthetic data generation scheme, see the supplementary materials in Pieciak et al. (2023).

We generate two sets of synthetic data representing a single fibre bundle (α_2_**=0**) and a two-fibre bundle case (α_1_**=**α_2_**=0.5**). The experimental setup for each dataset comprises 18 replicas of baseline samples and diffusion-weighted samples generated at ***b* = {600**, **1200} s /mm**^**2**^ and 90 gradient directions per shell distributed according to the sampling defined by the HCP WuMinn project (Van Essen, 2013). The signal-to-noise of the data ratio has been defined in terms of baseline signal as **SNR =*A*(0)/** *σ* and fixed to 300. For each set, we also generate noiseless reference data with no FW component, i.e., ***f* =0** in Eq. (1).

### 2.2. In vivo data

We use three publicly available diffusion-weighted MR databases compatible with single- and multiple-shell DTI and FW-corrected DTI. Two databases, namely MICRA (Koller et al., 2021) and Magdeburg (Lehmann et al., 2021), cover repeated scans for each subject acquired using a single scanner (i.e., intra-scanner acquisitions). The third database, the ZJU (Tong et al., 2020), covers multiple scans across identical scanners located in different centres (i.e., inter-scanner scenario).

The acquisition setup for the MICRA database was as follows:

- MICRA: The database covers inter-session repeated scans from a single centre. Six healthy volunteers (3F/3M) aged 24–30 were scanned, five times each using a 3T Connectom MRI research scanner (Siemens Healthcare, Erlangen, Germany) equipped with an ultra-strong gradient system at 300 mT/m. Acquisition protocol: single-shot spin-echo echo planar imaging sequence, anterior-posterior (AP) phase-encoding direction, repetition time (TR): 3000 ms, echo time (TE): 59 ms, pulse separation/pulse duration Δ/δ: 24/7 ms, field of view (FOV): 220 × 220 mm^2^, matrix size: 110 × 110, voxel size: 2 × 2 × 2 mm^3^, *b*-values: (200, 500, 1200, 2400, 4000, 6000) s/mm^2^ with (20, 20, 30, 61, 61, 61) gradient directions, respectively, 11 non-diffusion-weighted scans in AP direction repeated every twentieth volume and two non-diffusion-weighted scans in PA direction.

The acquisition details of two other databases are provided in the Supplementary materials. **Table 1** characterises all three databases in terms of subjects, the number of centres, sessions and scans, and the selected portion of diffusion-weighted MR data used to estimate the FWVF, DTI and FW-corrected DTI measures in a multiple-shell scenario. In the case of a single-shell scenario, we use only data acquired under the *b*-value annotated with the symbol “^⦰^”.

**Table 1.** The summary of diffusion-weighted MR databases considered in the study. The table provides *b*-values and the number of gradient directions used in our experiments.

| Database | Subjects | Centres | Sessions per centre | $b$-values [s/mm<sup>2</sup>] (gradients) | Non-diffusion-weighted replicas |
| --- | --- | --- | --- | --- | --- |
| MICRA | 6 (3F/3M) | 1 | 5 | 500 (20), 1200<sup>∘</sup> (30) | 11 AP, 2 PA |
| Magdeburg | 29 (24F/5M) | 1 | 2 | 1000<sup>∘</sup> (38), 2000 (76) | 14 AP, 9 PA |
| ZJU | 3 (2F/1M) | 10 | 1 | 1000<sup>∘</sup> (30), 2000 (30) | 6 |
| Multi $b$-value | 1 | 1 | 1 | 400 (33), 800 (33), 1000 (33), 1400 (33), 1600 (33), 2000 (33) | 1 |

<sup>∘</sup> Data acquired at annotated $b$ -value is used to obtain single-shell FWVF, DTI and FW-corrected DTI.

### 2.2 Data preprocessing

- MICRA: The dataset was preprocessed using the following pipeline: 1) noise removal using the Marčenko-Pastur Principal Component Analysis technique over window of size 5 × 5 × 5 voxels (MP-PCA; Veraart et al., 2016a; Veraart et al., 2016b), 2) Gibbs ringing artefacts correction (Kellner et al., 2016), 3) Rician bias correction applied voxel-wisely with the formula (Pieciak et al., 2018)

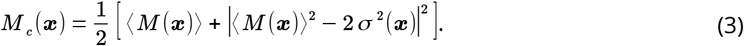

The correction given by Eq. (3) was applied under the assumption that the MP-PCA algorithm generates a proxy for the expectation of signal magnitude, **⟨ *M* (*x*)⟩**, *σ* ^2^(***x***) is the spatially-dependent noise standard deviation, 4) susceptibility-induced distortions estimation using the FSL FMRIB Software Library v6 `topup` tool (Analysis Group, FMRIB, Oxford, UK; Andersson et al., 2003; Smith et al., 2004), 5) head movements and eddy current distortions correction using the FSL `eddy` (Andersson et al., 2016), and 6) B1 field inhomogeneity correction using the N4 algorithm (Tustison et al., 2010).

### 2.4. Free-water-corrected Diffusion Tensor Imaging

The FW-corrected DTI is modelled using a two-component representation characterizing hindered diffusion using a diffusion tensor, and free diffusion represented by a mono-exponential decay (Pierpaoli and Jones, 2004; Pasternak et al., 2009):

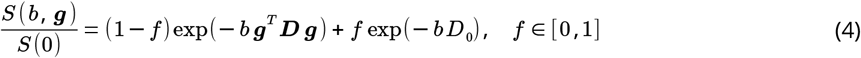

with ***S* (*b***, ***g*)** being the diffusion-weighted MR signal acquired in direction ***g*** at b-value ***b, S* (0)** is a non-diffusion-weighted MR signal, and ***D*** is a symmetric semi-positive matrix of size 3 × 3. The two-component representation given by Eq. (4) reduces to the standard DTI for ***f* = 0**.

### 2.5. Estimation methods: DTI and FW-corrected DTI

The following methods are considered in this study to estimate FWVF, DTI and FW-corrected DTI:

1. Bi-tensor-S: joint estimation of FWVF and FW-corrected DTI in the variational framework from single-shell acquisitions (Pasternak et al., 2009); the learning rate has been fixed to 0.005, the optimization uses 200 iterations and the initialization employs the tissue’s MD prior set to **0.6 × 10**^**−3**^ **mm**^**2**^**/ s** (Golub et al., 2020), unless otherwise stated,

2. Bi-tensor-M: joint estimation of FWVF and FW-corrected DTI *via* the non-linear least squares procedure from multiple-shell acquisitions (Hoy et al., 2014; Henriques et al., 2017),

3. SM: FWVF estimation using the spherical means technique from multiple-shell acquisitions (Tristán-Vega et al., 2022); the spherical harmonics decomposition at the order of ***L* = 6** is computed using the inverse linear problem with the regularization based on the Laplace-Beltrami operator and the regularization weight set to λ **= 0.001**, the parallel diffusivity parameter is fixed to λ_par_ = 2.0 × 10^−3^ mm^2^/ s and penalty term ν used to promote prolate convolution kernels have been optimized for each database separately: *ν* **= 0.015** (MICRA), *ν* **= 0.05** (Magdeburg), *ν* **= 0.05** (ZJU) and *ν* **= 0.05** (multi-*b*-value data).

4. DTI: standard DTI estimated *via* the non-linear least squares (NLLS; Koay et al., 2006),

5. FW-DTI: customized FW-corrected DTI scheme using the NLLS based on a pre-estimated FWVF with the SM technique. This scenario assumes the diffusion-weighted MR signal is corrected for the isotropic component prior to the estimation procedure:

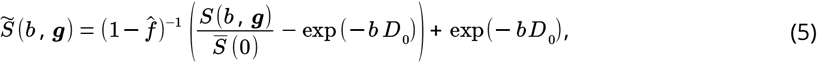

where 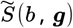 is the normalized diffusion-weighted MR signal after removing the FW component, 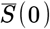 is the averaged signal across all non-diffusion-weighted MR acquisitions, 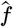 is the FWVF estimated using the SM technique from multiple-shell data.

We calculate FA, MD, RD and AD measures for each DTI-based technique considered in the study, i.e., Bi-tensor-S, Bi-tensor-M, DTI and FW-DTI. For DTI and FW-DTI, we handle two variants: single- and multiple-shell data. **Table 2** summarizes the methods used in the study.

**Table 2.** Summary of FWVF, DTI and FW-corrected DTI estimation procedures used in the study. The methods are classified into single- and multiple-shell data handled in the estimation process.

| Method | Single-shell |  |  | Multiple-shell |  |  |
| --- | --- | --- | --- | --- | --- | --- |
|  | FWVF | DTI measures | FW-corrected DTI measures | FWVF | DTI measures | FW-corrected DTI measures |
| **Bi-tensor-S** (Pasternak et al., 2009) | ✓ |  | ✓ |  |  |  |
| **Bi-tensor-M** (Hoy et al., 2014) |  |  |  | ✓ |  | ✓ |
| **SM** (Tristán-Vega et al., 2022) |  |  |  | ✓ |  |  |
| **DTI** (Koay et al., 2006) |  | ✓ |  |  | ✓ |  |
| **FW-DTI** |  |  | ✓ |  |  | ✓ |

We used DIPY library v. 1.9.0 (https://dipy.org) for non-linear DTI and DIPY with in-house implementation for FW-DTI fitting procedures delivered in Python 3.11.5 (https://www.python.org), NumPy 1.26.4 (https://numpy.org) and SciPy 1.11.1 (https://scipy.org). The implementation of Bi-tensor-S followed that of (Golub et al., 2021; https://github.com/mvgolub/FW-DTI-Beltrami), while Bi-tensor-M employed the one provided with DIPY library. The SM was estimated using the dMRI-Lab toolbox (https://www.lpi.tel.uva.es/dmrilab) run under MATLAB R2023b (The MathWorks, Inc., Natick, MA).

### 2.6. Data registration

The FA volumes estimated using a standard DTI at ***b* = 1200/ 1000 /1000 s / mm**^**2**^ for MICRA/Magdeburg/ZJU databases were used to register the data to the Montreal Neurological Institute (MNI) space, so-called standard space. Specifically, we linearly registered FA volumes computed with the standard DTI from each scan to the FSL template FMRIB58_FA using the FSL flirt tool under seven degrees of freedom, normalized correlation cost function and spline interpolation. The linearly transformed FA volumes were then non-linearly deformed *via* the FSL fnirt. All measures from the subjects’ native spaces were mapped to the MNI space using a trilinear interpolation. We then retrieved the white matter label from the Johns Hopkins University (JHU) WM atlas (Mori et al., 2005) in the standard space and shrunk it using a morphological binary erosion operator with a cross-shaped kernel of size 3 × 3 × 3 to eliminate potential misregistration outliers due to a partial volume effect. All scripting was carried out using the Arturo programming language 0.9.83 (https://arturo-lang.io).

### 2.7. Variability, separability and reliability assessment

Three characteristics of the FWVF and DTI-related measures are computed, namely reproducibility, reliability, and separability, all three in the standard space. First, we warp the measures from the subjects’ native spaces to the standard space using the warping fields obtained from the data registration procedure. Then, we compute the spatially dependent characteristics mentioned above across the repeated scans. Repeatability and reproducibility is explained in terms of the variability index, which defines the general ability to replicate the measure across scans or scanners of the same subject, and it is expressed by the coefficient of variation (CoV). By definition, the smaller the variability, the higher the repeatability and reproducibility. In our study, we distinguish three scenarios: 1) intra-scanner repeatability – the inter-sessions scans are replicated using the same scanner with a short time interval between the scans, 2) inter-session longitudinal reproducibility – the scans are repeated using the same scanner with a longer time interval between the scans (e.g., several weeks) and possibly affected by confounding factors (e.g., changes in magnetic field drifts) and 3) inter-scanner reproducibility – the scans are repeated using the same scanner type, but installed in different locations. The reliability index reflects the measure’s consistency, and it is expected to be high for measures that are stable in value across repeated scans. Decomposing the variance into within-subject and between-subjects variances, as defined by Zuo et al. (2019), enables the computation of a ratio that reflects how much of the total variance is explainable by actual inter-individual differences rather than noise or session variability. The last index, separability, illustrates the metric’s ability to capture inter-subject discrepancies. Before introducing these indices, we define the statistics computed across the measures in the standard space.

#### Statistics across the scans

Given the subject ***s* = 1**, **…, *S***, we define the sample mean and sample standard deviation across the stack of measures as follows

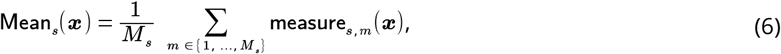

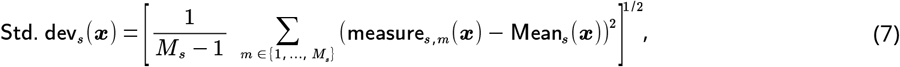

where **measure**_*s, m*_ **(*x*)** is the measure computed for the ***s* -th** subject and the ***m* -th** scan with *m* = 1, …, *M*_s_. The Eq. (7) reduces to **Std. dev_s_ (*x*) = 2**^**−1/2**^ **×** |**measure_s,1_ (*x*) − measure_s,2_ (*x*)**| for a test-retest case. The statistics given by Eqs. (6) and (7) are spatially dependent, i.e., they are computed for each spatial location ***x*** in the standard space.

#### Variability

The variability of the metric has been defined as the median value from CoVs across all subjects

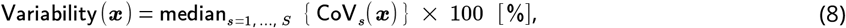

where the coefficient of variation **CoV**_***s***_**(*x*)** for subject ***s*** is given by

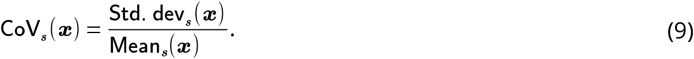

In the case of two samples (i.e., test-retest data), the formula (9) reduces to:

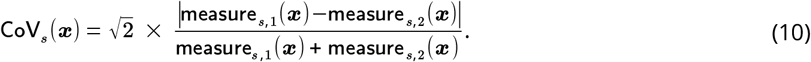

#### Reliability

The reliability index has been defined using the formulation by Zuo et al. (2019)

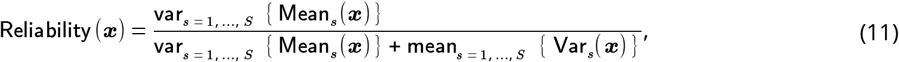

where **Var** _*s*_**(*x*)** is the sample variance of the measure calculated across the scans for ***s* -th** subject and **var**_*s* = 1, …, *S*_ **{.}** is the population variance calculated across sample means **Mean**_*s*_**(*x*)**, i.e., the normalization factor in the variance formula equals **1 /*S***.

#### Separability

We define the separability index as follows:

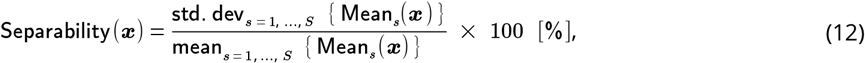

where **std. dev**_*s* = 1, …, *S*_ **{**. **}** is the population standard deviation with the normalization factor also given by **1 /*S***.

#### Diffusion kernel density estimation (DiffKDE)^2^

The DiffKDE method estimates the probability density function from data samples in a non-parametric way. Given the samples ***x***_**1**_,**…, *x***_***n***_ from a probability density function ***f* (*x*)**, kernel *K* and bandwidth ***h* >0**, the formula for the KDE at a point ***x***_**0**_ is given by (Hastie et al., 2009)

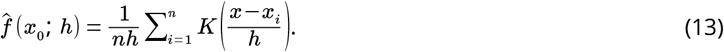

To determine the bandwidth *h*, we use the improved Sheather-Jones algorithm, which minimizes the asymptotic mean squared error between true probability density function ***f* (*x*)** and the estimated 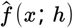 (Sheather and Jones, 1991; Botev et al., 2010)

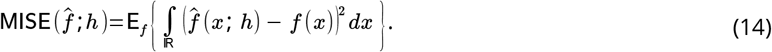

The statistics above-presented will be computed in the following three scenarios:

1. inter-session MICRA database (***S* = 6**, *M* _*s*_ = 5),

2. inter-session longitudinal Magdeburg database (***S* = 29, *M*** _*s*_ **= 2**),

3. inter-scanner ZJU database (***S* = 3**, *M* _*s*_ = 10).

## 3. Experimental results

This section wraps the experimental results generated for *in silico* data and MICRA database. The results for the Magdeburg and ZJU databases are included in the supplementary materials, and if necessary, references to specific results are given in this section.

### 3.1. In silico experiments

In the first *in silico* experiment depicted in **Fig. 1**, we examine different FW-correction DTI schemes and relate the FW-corrected MD and FA measures under ***f* =0.2** to the standard DTI-based equivalents, but obtained from the cellular model only (i.e., ***f* =0** in Eq. (1)). Among the multiple-shell-based techniques, the FW-DTI customised scheme yields smaller discrepancies of MD and FA from the standard DTI-based MD and FA than the B-tensor-M technique. However, the FW-corrected MD using the FW-DTI scheme is more flattened compared to the FW-corrected MD obtained *via* the Bi-tensor-M. Regarding the single-shell techniques evaluated at ***b* = 1200 s/ mm**^**2**^, the flattening effect is more evident, albeit the FW-corrected MD using the FW-DTI scheme also follows the general trend of standard DTI-based MD. The Bi-tensor-S method obtains the tissue’s prior, i.e., the FW-corrected parameter oscillates around the value of **0.6 × 10**^**−3**^ **mm**^**2**^**/ s**. The extended experiment presenting the estimated tissue’s prior using the Bi-tensor-S method is shown in **Fig. S2**. These *in silico* experimental results are consistent between single- and two-fibre bundle scenarios (cf. **Fig. 1** to **Fig. S1)**.

**Figure 1.**
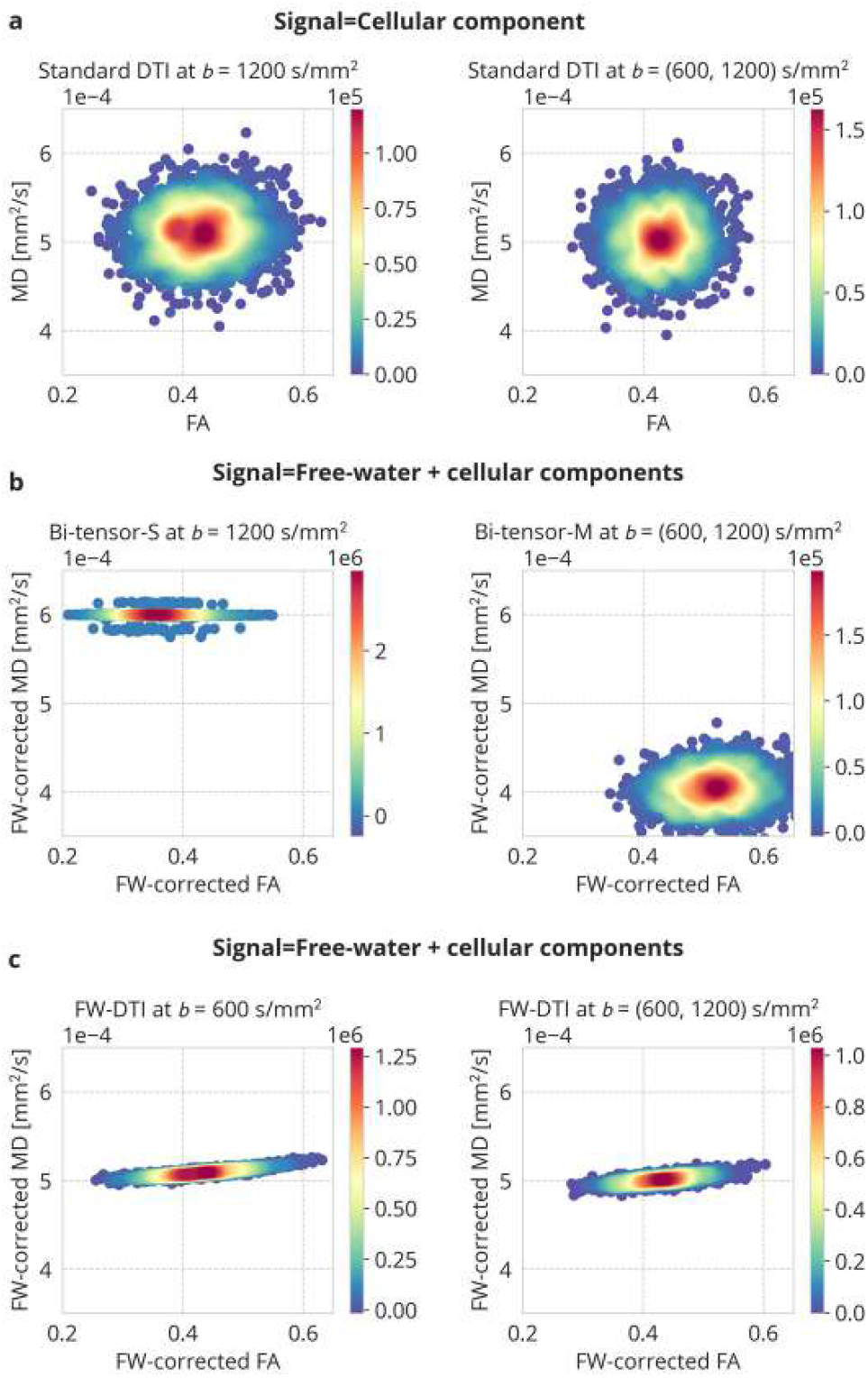
The 2D density plots illustrate the experimental results from an *in silico* model under a two-fibre bundle configuration: **a)** the MD and FA parameters estimated using the standard DTI from noiseless reference data covering the cellular component only. The FW-corrected MD and FA estimated from the signal covering FW (***f* =0.2**) and cellular components: **b)** Bi-tensor-S and Bi-tensor-M approaches, and **c)** FW-DTI customised scheme according to Eq. (5). In total, 15^3^ samples have been used to obtain a single 2D density plot.

The second experiment, as illustrated in **Fig. 2**, quantitatively examines the FW-corrected FA and MD measures as a function of ground-truth FWVF parameter, and verifies the precision of the FWVF parameter estimated using different procedures considered in the study. The misestimation of the FW-corrected DTI-based measures is quantified using the mean percentage error, i.e., the mean absolute value of the relative error, and the standard deviation of the percentage error. We observe that the FW-DTI scheme is characterized by the smallest mean error among the evaluated methods, with a mean percentage value fluctuating between 2.75% and 6% depending on the measure and experimental variant, i.e., single- or multiple-shell data. The SM technique estimates the FWVF parameter most accurately, with almost no error dependence on the reference FWVF value, as observed particularly in the case of the Bi-tensor-S and Bi-tensor-M techniques. However, we observe an increased standard deviation for the SM technique compared to the Bi-tensor-M, but it is still smaller than that of the Bi-tensor-S.

**Figure 2.**
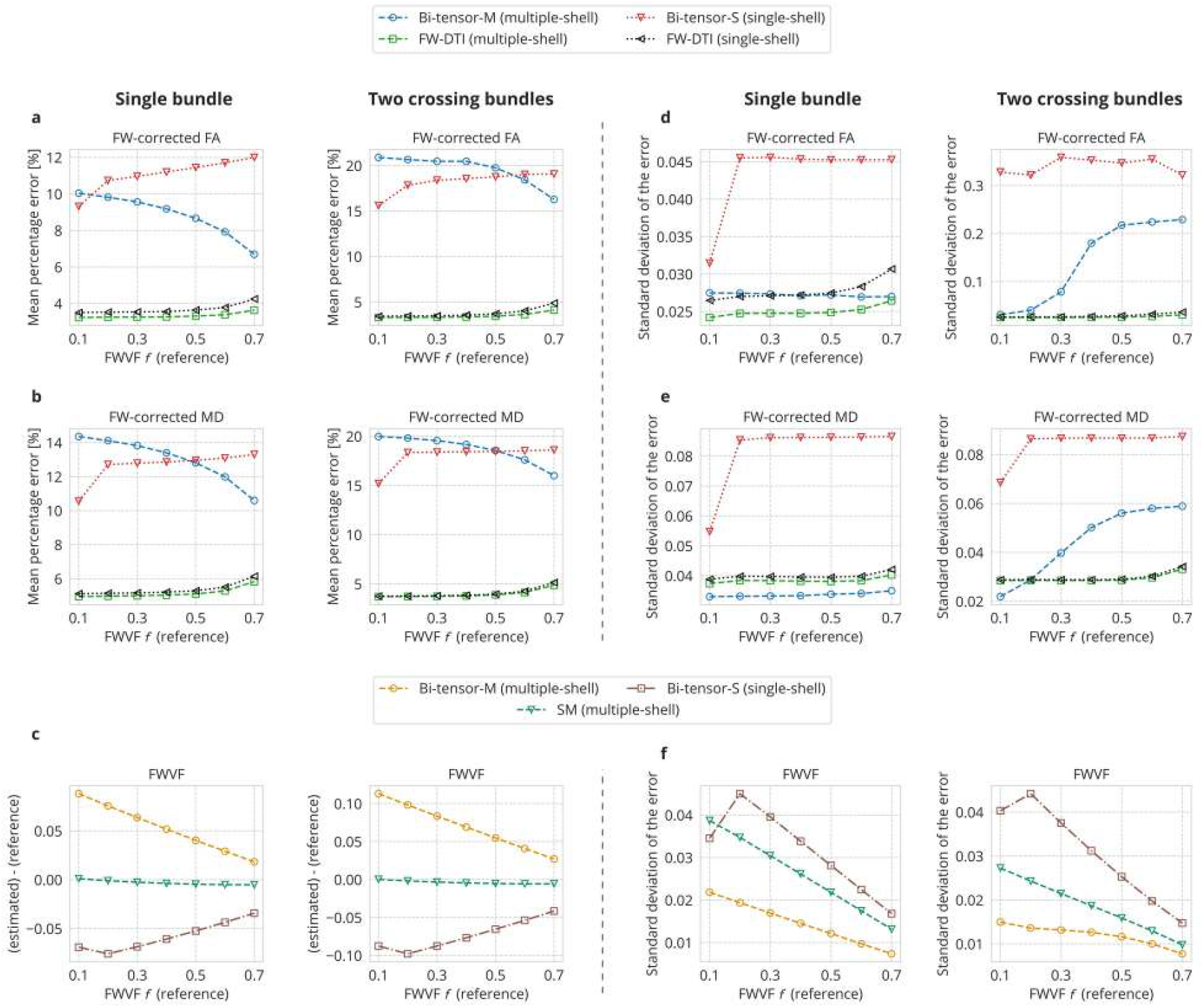
The mean percentage error and standard deviation of percentage error of DTI-based measures estimated from an *in silico* model: **a**,**d)** FW-corrected FA and **b**,**e)** FW-corrected MD as a function of reference FWVF ***f***, depending on the methodology used to correct the DTI for the FW. **c)** The mean difference between the estimated FWVF and reference FWVF and **f)** the standard deviation computed from the differences between the estimated FWVF and reference FWVF as a function of reference FWVF. The left column in each panel presents the results for a single-fibre bundle configuration, while the right illustrates a two-fibre bundle configuration in the model defined by Eq. (5). In total, 15^3^ samples have been used to obtain a single value in each plot.

### 3.2 Visual inspection of the measures

We now move to *in vivo* experiments, first by visually inspecting the measures in the subject’s native coordinate system. We have chosen a single subject from the MICRA database (sub-01, ses-01) and displayed the measures in **Fig. 3**. The results for the inter-session longitudinal Magdeburg database and inter-scanner ZJU database are included in the Supplementary materials as **Fig. S3** and **Fig. S4**, respectively. In **Fig. 3a**, we illustrate the FWVF estimated under multiple-shell data at ***b* = {500**, **1200} s /mm**^**2**^ using the SM and Bi-tensor-M approaches, and single-shell data at ***b* = 1200 s/ mm**^**2**^ with the Bi-tensor-S. The results obtained with Bi-tensor-M and Bi-tensor-S differ despite both techniques are conceptually based on a direct optimization of Eq. (4), albeit using distinct numerical schemes. Next, in **Fig. 3b** and **Fig. 3c**, the DTI measures estimated from multiple- and single-shell acquisitions have been illustrated in the following rows: i) standard DTI (no free-water correction), ii) FW-corrected DTI according to Eq. (5) and iii) Bi-tensor-M or Bi-tensor-S approach. We observe increased FA values and decreased MD, AD and RD over the brain using all three FW-correction schemes examined, i.e., the FW-DTI, Bi-tensor-M and Bi-tensor-S. Besides, the FW-corrected MD parameter estimated using the Bi-tensor-S approach exhibits flattened spatial characteristics over the brain (see **Fig. 3c**). The median (interquartile range) value of FW-corrected MD computed with the Bi-tensor-S approach over the WM area is **0.6006 (0.6 – 0.6015)**. This characteristic of the Bi-tensor-S-related MD parameter has been observed in three considered databases (see also **Fig. S3c** and **Fig. S4c**).

**Figure 3.**
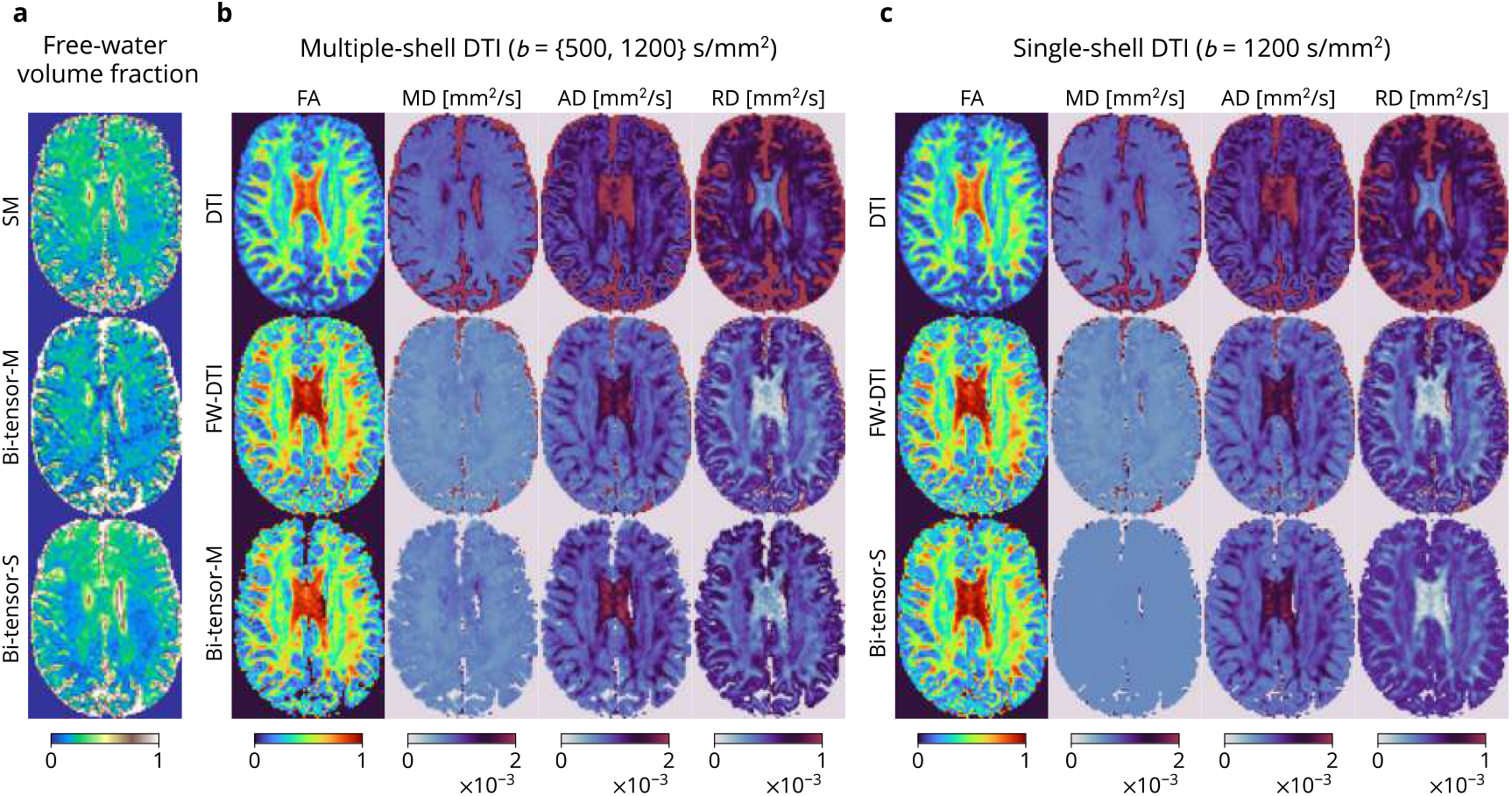
Estimated microstructural measures for a selected MICRA acquisition (sub-01, ses-01, slice 36): **a)** FWVF estimated from multiple-shell data (SM and Bi-tensor-M approaches) and single-shell data (Bi-tensor-S), **b)** DTI-based measures estimated from multiple-shell data using a standard DTI, and two free water-correction methodologies: FW-DTI according to Eq. (5) and Bi-tensor-M, and **c)** DTI-based measures estimated from single-shell data using the standard DTI, FW-DTI and Bi-tensor-S. The FWVF parameter for FW-DTI was pre-estimated using the SM approach from multiple-shell data in both scenarios presented in panels b) and c).

### 3.3. Variability maps

In the experiment shown in **Fig. 4**, we visually inspect spatially-dependent variability maps of the measures in the standard space computed according to Eq. (8) for the MICRA dataset. The results for two other datasets have been included in the Supplementary materials, specifically in **Fig. S5** (Magdeburg) and **Fig. S6** (ZJU).

**Figure 4.**
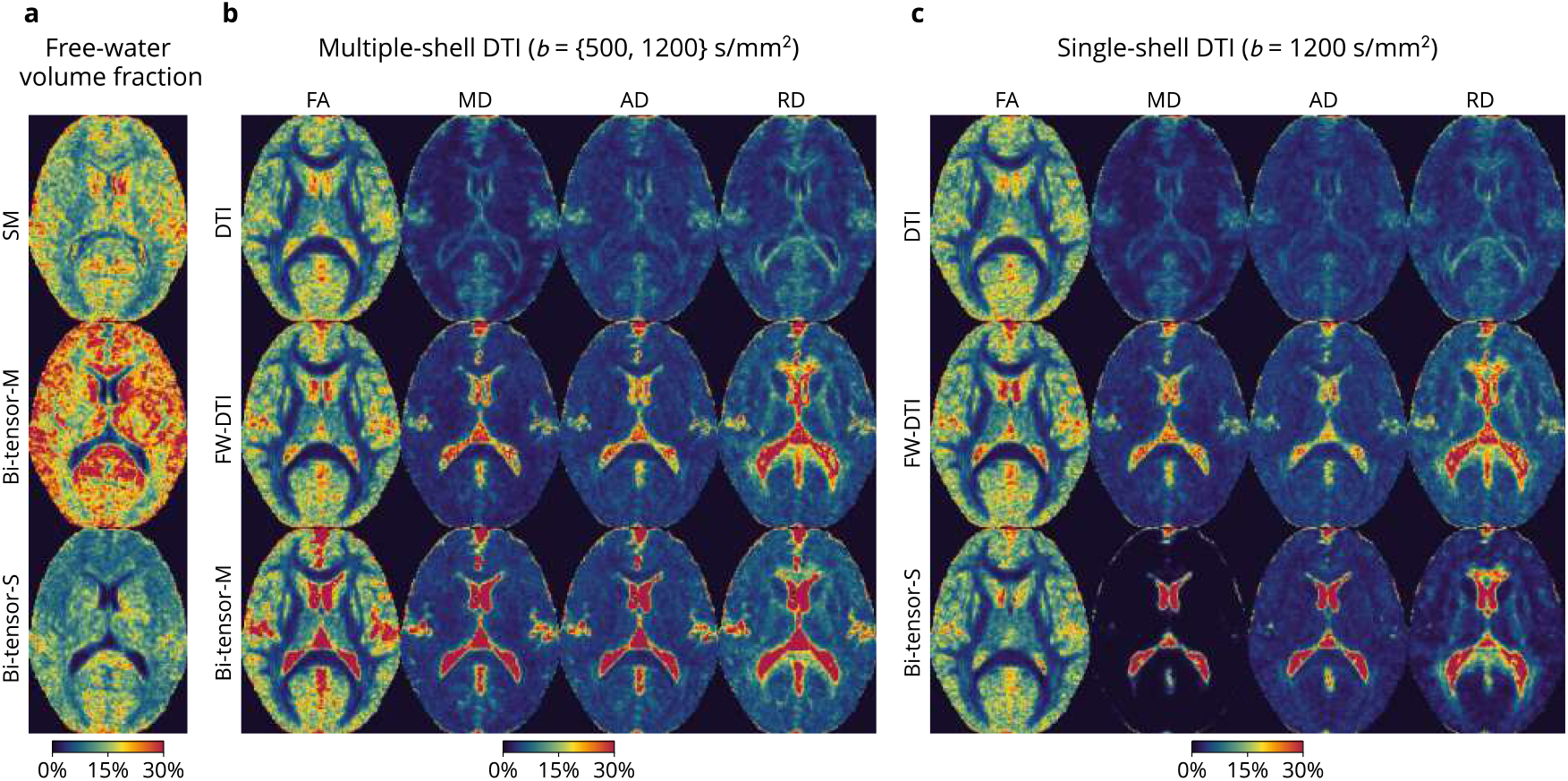
Inter-session variability maps of the measures defined in the standard space for MICRA database according to the coefficient of variation defined by Eq. (8): **a)** FWVF estimated from multiple-shell data (SM and Bi-tensor-M approaches) and single-shell data (Bi-tensor-S), **b)** DTI-based measures estimated from multiple-shell data using a standard DTI, and two FW-correction methodologies: FW-DTI according to Eq. (5) and Bi-tensor-M, and **c)** DTI-based measures estimated from single-shell data with the standard DTI, FW-DTI and Bi-tensor-S. The FWVF parameter for FW-DTI was pre-estimated using the SM approach from multiple-shell data in both scenarios presented in panels b) and c).

In **Fig. 4a**, we illustrate the variability of the FWVF estimated from: i) multiple-shell data using the SM, ii) multiple-shell data using the Bi-tensor-M procedure and iii) single-shell data *via* the Bi-tensor-S. Among the three techniques mentioned above, we observe the highest variability of the FWVF parameter computed with the Bi-tensor-M approach. However, the variability of the FWVF deviate across the datasets examined in the study, and tends to be visually smaller in the case of Bi-tensor-M technique for Magdeburg and ZJU databases than those of other techniques (cf. **Fig. 4a** to **Fig. S5a** and **Fig. S6a)**.

Next, in **Fig. 4b** and **Fig. 4c**, we present the variability of DTI metrics computed with the standard DTI and under a FW-correction from multiple- and single-shell data. The first observation is that the variability of the FA parameter follows different spatial characteristics compared to MD, AD and RD. Specifically, the variability of FA is remarkably lower in the white matter compared to the grey matter, while the variability of MD, AD, and RD is more homogeneous across the white and grey matter areas. However, in the case of MD estimated with Bi-tensor-S (see also **Fig. S6c**), the results are contentious – the experiment has revealed the smallest variability across all the measures considered in the study. As we will demonstrate later, the reliability of the FW-corrected MD parameter, particularly when estimated using the Bi-tensor-S approach, is low compared to other metrics.

### 3.4. Density-based variability and reliability indices

The following two experiments assess the variability and reliability of the measures across the white matter in the form of density plots. The plots have been estimated using the DiffKDE approach (Sheather and Jones, 1991; Botev et al., 2010) for all measures previously illustrated in **Fig. 3** and are depicted in **Fig. 5** (MICRA). The results for two other datasets are included in the Supplementary materials as **Fig. S7** (Magdeburg) and **Fig. S8** (ZJU).

**Figure 5.**
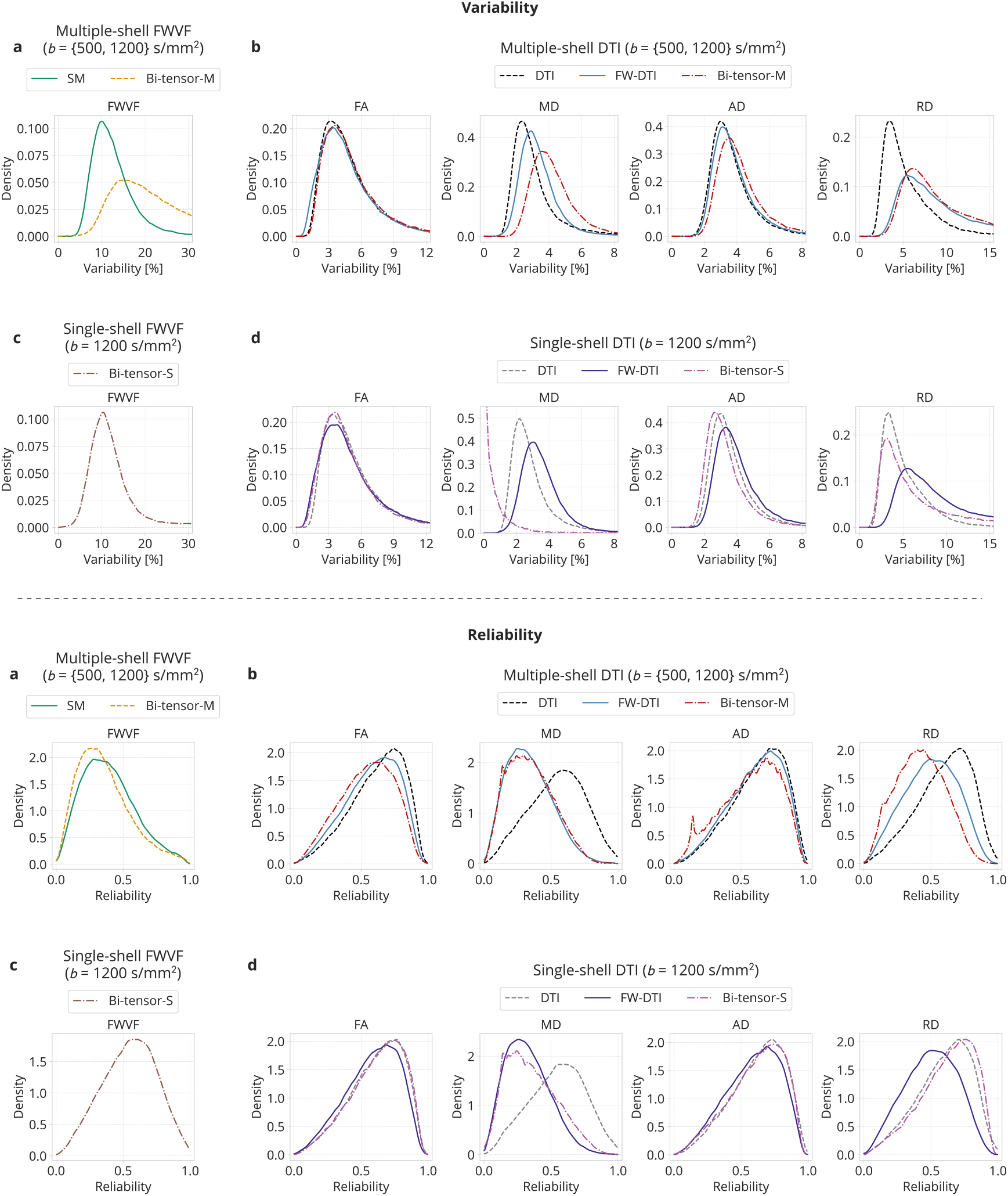
Inter-session kernel density-based variability (top) and reliability (bottom) indices computed for MICRA database over the white matter area: **a)** FWVF estimated from multiple-shell data (SM and Bi-tensor-M approaches), **b)** DTI-based measures estimated from multiple-shell data using standard DTI, FW-DTI according to Eq. (5) and Bi-tensor-M, **c)** FWVF estimated from single-shell data (Bi-tensor-S) and **d)** DTI-based measures estimated from single-shell data with the standard DTI, FW-DTI and Bi-tensor-S. The FWVF parameter for FW-DTI was pre-estimated from multiple-shell data using the SM approach in both scenarios presented in panels b) and d).

The FWVF parameter, no matter how it is estimated and in which variant (single- or multiple-shell data), is poorly reproducible over the white matter, to a much lesser extent than the DTI and FW-corrected DTI measures. Density plots representing the variability (i.e., CoV parameter) of the FWVF parameter reach their peaks at **CoV=10.08 %** (SM), **CoV=15 %** (Bi-tensor-M) and **CoV=10.31 %** (Bi-tensor-S). In general, the plots representing the variability for all measures considered in the study exhibit unimodal, predominantly positively skewed densities. Our results demonstrate the FW-correction primarily does not improve the repeatability/reproducibility of DTI-related measures compared to the standard DTI case, and even if it does, the changes are minute, mainly observed in the Bi-tensor-S approach (see **Fig. 5d** and **Fig. S8d**). We found the contrary behaviour of multiple-shell Bi-tensor-M scheme – the FW-correction has led to an increase in the variability of MD/AD/RD measures, the results being unfailingly observed in all three experimented databases (**Fig. 5c, Fig. S7c** and **Fig. S8c**).

Our experiments also reveal that the measures characterised with the lowest reliability are multiple-shell FWVF and FW-corrected MD obtained from either single-or multiple-shell data (see **Fig. 5**). Notably, the FW-correction to MD has led to a significant decrease in the reliability parameter, regardless of the numerical method used to estimate the measure and the data type handled, i.e., the reliability peak is far below 0.5. At large, the plots representing the reliability index exhibit unimodal and negatively skewed densities for the evaluated measures (cf. to the densities of the variability parameter). In general, the reliability of standard DTI-based measures is roughly equal to or better than the reliability of equivalent measures corrected for the FW component. These results are compatible between all three databases considered in the study.

### 3.5. Variability, reliability and separability

We now proceed to quantitative experiments that demonstrate two relationships: i) reliability versus variability and ii) separability versus variability.

In the first experiment depicted in **Fig. 6**, we put together the population’s first (25th percentile), second (median) and third (75th percentile) quartiles computed for the variability and reliability indices. The “population” is understood here as all voxels taken from the white matter region defined in the standard space. Note that the smaller the third quartile for the variability index, the better, while the larger the third quartile for the reliability, the better. As mentioned in the previous section, the FWVF is characterized by the highest variability among all measures, typically several times higher than DTI-based measures, and a relatively low reliability. However, the variability and reliability indices vary between the methods used to estimate the FWVF and across the datasets (cf. **Fig. 6a**,**c** to **Fig. S9a**,**c** and **Fig. S10a**,**c**). Nevertheless, the SM technique has shown superior behaviour over the Bi-tensor-M in terms of reliability in all three databases tested, though the variability index does not give a clear answer to which method varies more. The multiple-shell DTI-based measures with no FW correction are more reproducible and more reliable than FW-corrected equivalents using the Bi-tensor-M approach. The experiments show evidence that correcting DTI measures for the FW component *via* Eq. (5) would be a more appropriate solution, given that it potentially avoids such declines in reproducibility and reliability indexes, as observed in the Bi-tensor-M approach.

**Figure 6.**
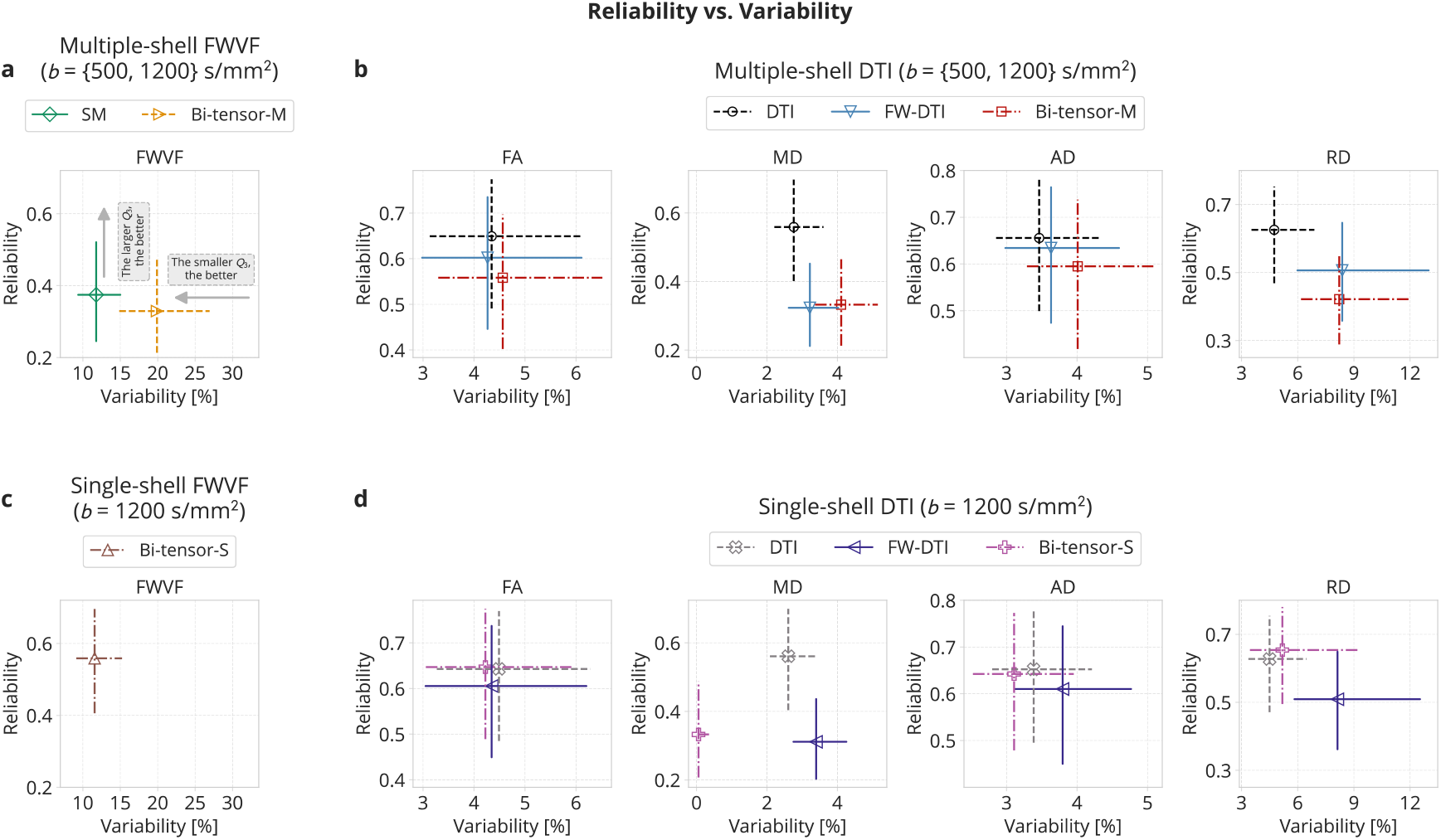
Inter-session reliability versus variability plots computed for MICRA database over the white matter area: **a)** FWVF estimated from multiple-shell data (SM and Bi-tensor-M approaches), **b)** DTI-based measures estimated from multiple-shell data using standard DTI, FW-DTI according to Eq. (5) and Bi-tensor-M, **c)** FWVF estimated from single-shell data (Bi-tensor-S) and **d)** DTI-based measures estimated from single-shell data with the standard DTI, FW-DTI and Bi-tensor-S. The FWVF parameter for FW-DTI was pre-estimated from multiple-shell data using the SM approach in both scenarios presented in panels b) and d). The markers present median values calculated over the white matter area, while the horizontal and vertical lines represent distances between the first ***Q***_**1**_ **= 0.25** (25th percentile) and third ***Q***_**3**_ **= 0.75** (75th percentile) quartiles. The horizontal and vertical ranges for plots representing multiple- and single-shell equivalent cases have been fixed.

The results obtained in a single-shell scenario *via* the Bi-tensor-S are generally inconsistent across the datasets employed in the study. For instance, the FW-corrected AD and RD parameters seem to be more reproducible and somewhat more reliable than the DTI-based ones for the ZJU database (inter-scanner acquisitions), while in the case of the Magdeburg database (longitudinal intra-scanner), we observe the opposite trend (cf. **Fig. S9d** to **Fig. S10d**). These inconsistencies are also observed for the variability index between two inter-session databases, i.e., MICRA and Magdeburg. Specifically, the MD parameter estimated with Bi-tensor-S exhibit the smallest and the largest variability among single-shell-based methods, respectively (cf. **Fig. 6d** and **Fig. S10d** to **Fig. S9d**).

In the final experiment, we strive to establish optimal measures in terms of reliability, variability and separability indices. In the bar charts presented in **Fig. 7a**, we relate the median reliability to median variability. Both indices are ordered according to the median reliability. The indices were computed over the white matter area in the standard space. In general, the results presented in charts-based plots illustrate that the highest reliability measures are generally approvingly reproducible, with the variability being less than 5% for inter-session and inter-session longitudinal acquisitions (see also **Fig. S11a**) and oscillating around 5% for inter-scanner acquisitions (**Fig. S12a**). In this class, one can identify AD, FA, and RD computed from standard DTI, with the last two measures characterized by higher variability and separability than the AD (**Fig. 7b**). Contrarily, the measures characterized by low reliability, such as the FWVF, are typically highly variable.

**Figure 7.**
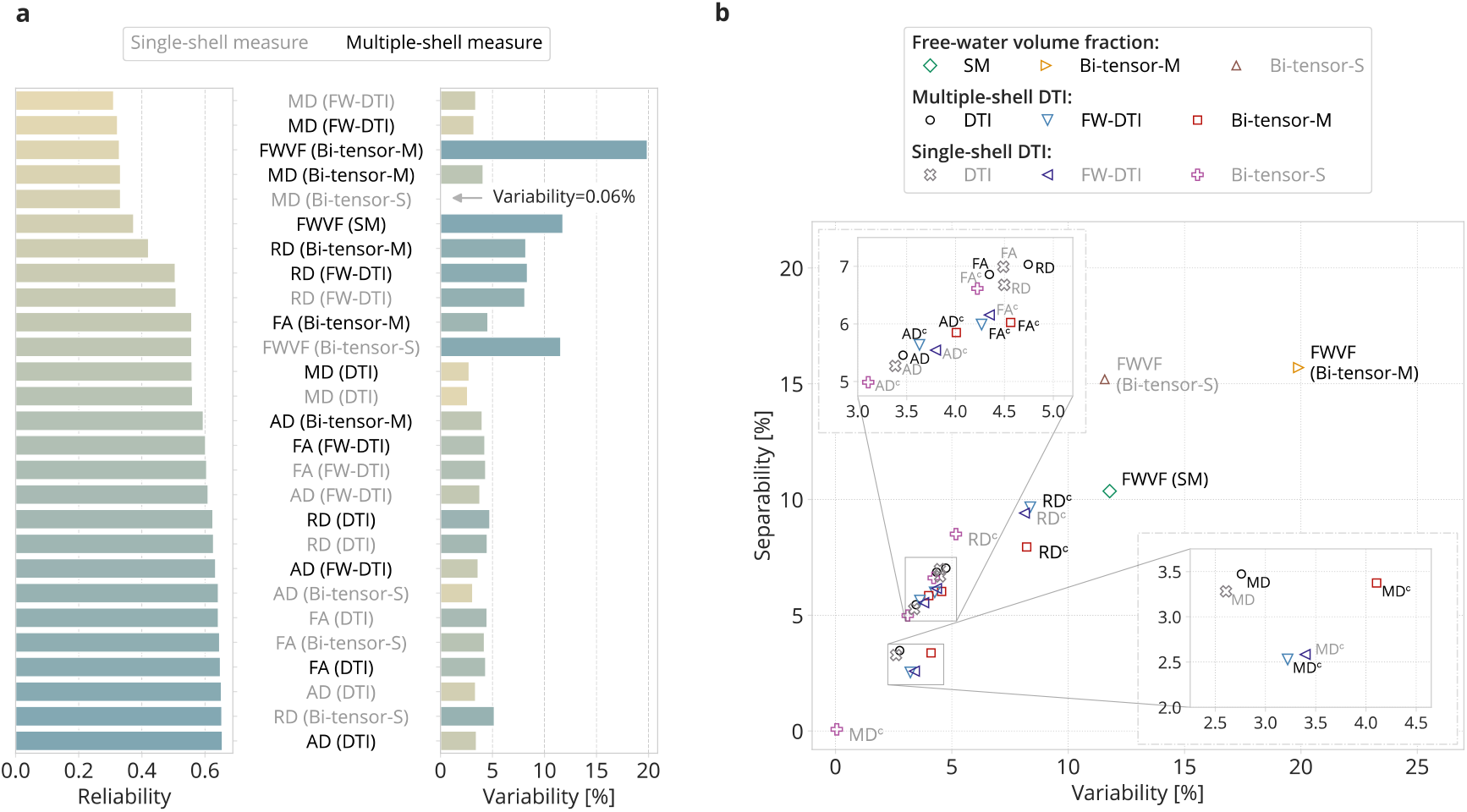
**a)** Box-plots presenting inter-session median reliability and median variability indices computed from the MICRA database over the white matter area. The colours in the variability box-plot reflect the ordering according to the reliability index value. **b)** A diagram representing the separability index as a function of the variability index. A single marker refers to the median variability and median separability values calculated over the white matter area in the standard space. Multiple-shell measures are annotated with black text, while the single-shell measures are in grey. The FW-corrected DTI measures are annotated with a subscript “c”.

Considering the FW-corrected DTI, we observe FA and AD rectified with the customized scheme from multiple-shell data are the most reliable among the FW-corrected DTI schemes, reproducible and somewhat separable across the FW correction schemes considered in the study. The previously discussed FW-corrected MD with the Bi-tensor-S, although it has revealed extremely low variability, is a non-reliable measure. Our experiments have also demonstrated that the FW-corrected MD under two other schemes, i.e., Bi-tensor-M and FW-DTI, are also the least reliable among the DTI measures, and this result is compatible across the databases considered in the study. Overall, the MD measure, computed in any FW-correction variant is the least reliable and separable parameter among all parameters considered in this study. The FWVF is also non-reliable but seems to be a highly separable parameter. Interestingly, the FW-corrected RD obtained with any FW correction scheme is moderately reliable, but reasonably highly variable and separable measure, second only to the FWVF.

## 4. Discussion

One can follow several approaches to correct the DTI for the FW, either using single- or multiple-shell diffusion-weighted MR data. The first and most popularized approach by Pasternak et al. (2009), which we refer to here as the Bi-tensor-S, directly optimizes the bi-tensor representation given by Eq. (4) within a variational framework. This formulation enables estimation of both the FWVF and FW-corrected DTI measures in a joint optimisation procedure using only single-shell diffusion-weighted MR data acquired approximately at ***b* = 1000 s/ mm**^**2**^. However, the method requires the initialization scheme to be carefully selected (Parker et al., 2020; Golub et al., 2021). The second group of methods optimize Eq. (4) using the numerical schemes tailored for multiple-shell acquisitions (Pasternak et al., 2012; Hoy et al., 2014; Bergmann et al., 2020). As a representative, in this study, we follow the region contraction-based method by Hoy et al. (2014), which we call the Bi-tensor-M. Recently, the advantage of FWVF estimated using multiple-shell over single-shell has been demonstrated in the context of brain age estimation (Nemmi et al., 2022) and healthy brain ageing (Correia et al. 2024). An alternative solution is to estimate the FWVF parameter, correct the diffusion-weighted MR signal for the FW component and then re-estimate the standard DTI from the corrected signal (Pieciak et al., 2023; Chang et al., 2024; Guadilla et al., 2025). The last group consists of deep learning-based approaches that aim to find a non-linear mapping between diffusion-weighted MR signal and FW or FW-corrected DTI parameters (Molina-Romero et al., 2018; Weninger et al., 2020).

We start our discussion by commenting on the sanity checks, displaying the estimated measures using different techniques. **Fig. 3d, Fig. S3d** and **Fig. S4d** depict particularly flattened FW-corrected MD characteristics computed with the Bi-tensor-S approach. The median [interquartile range] of FW-corrected MD measure over the WM area for two other databases is **0.5991 (0.5699 – 0.6285)** (Magdeburg) and **0.601 (0.5996 – 0.6034)** (ZJU). These numbers (along with the numbers obtained from MICRA database) correspond with those results presented in (Golub et al., 2021, Table 2). We note the flattening effect has been previously explored by Golub et al. (2021) in the context of *in silico* experiments and by Correia et al. (2024) in brain ageing. To put it differently, the FW-corrected MD using the Bi-tensor-S turns out to be the tissue’s prior (see **Fig. S2** for an extended *in silico* experiment). Contrary to Bi-tensor-S, the FW-corrected MD computed using the Bi-tensor-M and FW-DTI approaches has enabled to discriminate between WM and GM areas. As for other FW-corrected measures, increased FA and decreased MD/AD/RD parameters over the white matter are consistent across the datasets considered in our study and with previous reports (Metzler-Baddeley et al., 2012; Hoy et al., 2014; Golub et al., 2021; Pieciak et al., 2023).

It is noteworthy that the FWVF estimated using SM, Bi-tensor-S, and Bi-tensor-M actually presents the effective FW fraction confounded by *T*_*2*_ relaxation, but it is typically used as a proxy for the FWVF (Pasternak et al., 2009; Golub et al., 2021). Although none of the above-mentioned methods directly models freely diffusing water, they somewhat aggregate the diffusion found as the cerebrospinal fluid and interstitial fluid in the extracellular space of grey and white matter (Pasternak et al., 2009). Intrinsically, the FWVF may be biased by other pools, such as blood perfusion, which affects the signal at low *b*-values (Rydhög et al., 2018). **Fig. S13** illustrates the relationship between the FWVF parameter computed using the SM and Bi-tensor-M from the same multiple-*b*-value *in vivo* acquisition, considering different *b*-values configuration. This experiment gives evidence that the Bi-tensor-M approach provides smaller FWVF values than the SM technique for low *b*-values configurations (here, ***b* = {400, 1000} s / mm**^**2**^), while for higher *b*-values combinations, such as ***b* = {1000**, **2000 } s/ mm**^**2**^, we observe the opposite demeanour, i.e., Bi-tensor-M provides higher FWVF values than the SM. These results also reveal that the SM technique is more stable in the estimated values as the *b*-value changes. The results from this experiment are consistent with those variations we observe with the MICRA, Magdeburg and ZJU in **Fig. 3, Fig. S3**, and **Fig. S4**, respectively. Moreover, the SM technique, by its nature, enables handling complex geometries (e.g., crossing fibres) by modelling the signal with a continuous mixture of tensors in the space of orientations, thus reducing the systematic bias observed with the Bi-tensor-M approach.

In the study by Albi et al. (2017), it has been suggested that the FW correction to DTI from a single-shell acquisition improves the test-retest repeatability of FA and MD metrics by reducing the CoV *on average* approximately at 1%pt. Our study partially corroborates these results, illustrating an improvement in FA/MD repeatability for MICRA and reproducibility for ZJU databases in terms of median CoV over the WM at 0.3%pt./2.5%pt. and 1.1%pt./4.0%pt., respectively. However, this pattern is not regular, as our results for the inter-scanner longitudinal Magdeburg database manifest the opposite trend (-0.1%pt./-0.8%pt.; see also **Fig. S7d**). The recent study by Correia et al. (2024) discovered the flattening effect of FW-corrected MD parameter with age. Our study has revealed an excellent reproducibility of FW-corrected MD, which directly explains the flattened spatial characteristics of the measure (cf. the Bi-tensor-S results demonstrated in **Fig. 4c** to **Fig. 3d**). In other words, the Bi-tensor-S provides the prior for the MD measure (here, assumed to be **0.6 × 10**^**−3**^ **mm**^**2**^**/ s**) rather than the value contemplating the actual FW-corrected MD parameter. Moreover, the FW-corrected MD parameter with the Bi-tensor-S approach is neither reliable (see Fig. **Fig. 7a**) nor separable (Fig. **Fig. 7b**). This is a consequential result that may raise questions about the repeatability (and thus, the trustworthiness) of previous findings in the brain studies based on a single-shell Bi-tensor-S-based MD parameter. In general, the FW-corrected MD parameter estimated using any single-or multiple-shell-based method considered in our experiments exhibits low reliability and separability, which clarifies the factiously excellent reproducibility observed in **Fig. 4**. Interestingly, the reliability of FW-corrected MD measure with any method is consistently much less reliable than the equivalent standard DTI (see **Fig. 6b,d**). The results obtained in single-shell FW-corrected measures do not translate to a multiple-shell scenario, as the FW-corrected MD, AD and RD parameters computed with Bi-tensor-M reveal increased variability compared to the standard DTI (see **Fig. 6b** and **Fig. S10b**). A direct reason for the variability growth observed in the Bi-tensor-M approach is the increased number of degrees of freedom in the optimised cost function. The variability of a FW-corrected DTI might decrease if one pre-estimates the FWVF using an external method, such as the SM (Tristán-Vega et al., 2022), corrects the diffusion-weighted MR signal for the FW component and then re-estimates the DTI using a standard procedure. Remarkably, “fixing” the FWVF in the optimization process does not reduce the reliability of the FW-corrected measures compared to the Bi-tensor-M, as indicated by the experiment depicted in **Fig. 6b** (see also **Fig. S9b** and **Fig. S10b**).

The experiments demonstrated consistency in the variability of FWVF out against the DTI and FW-corrected DTI measures across all three datasets. Specifically, the FWVF estimated using any method considered in the study demonstrates higher CoV over the white matter area compared to all DTI-based measures examined in this study. The experimental results do not provide a clear answer as to which multiple-shell-based technique (i.e., the region contraction-based or the spherical means) is superior in terms of reproducibility. However, the Bi-tensor-M technique is trailing behind the other two methods (i.e., SM and Bi-tensor-S), considering the reliability index.

We note the study goes beyond the standard intra-site repeatability or inter-site reproducibility, as it also explores longitudinal reproducibility. Such longitudinal reproducibility evaluation is particularly important in a clinical scenario once the features observed in the images are expected to demonstrate the evolution of the brain between the scans, or confounding factors such as different operators handling the scanner or magnetic field drifts (Lehmann et al., 2021; Boudreau et al., 2025).

In our study, we have resorted to computing the KDE over the white matter area rather than reporting numbers reflecting the variability or reliability, as presented in other studies (Grech-Sollars et al., 2015; Albi et al., 2017). Note that, in general, potential misregistered voxels or the voxels representing the partial volume effect do not seriously affect the shapes of density plots. However, the KDE-based procedure must be applied with extreme care, as the standard Silverman’s bandwidth selection method (Silverman, 1986) is prone to errors for non-Gaussian histogram shapes. In this study, we have employed an advanced DiffKDE method with the improved Sheather-Jones algorithm for bandwidth selection (Sheather and Jones, 1991; Botev et al., 2010). This technique has enabled us to accurately reflect density plots, regardless of their shapes, as demonstrated in **Fig. S14**.

## 5. Conclusions

This paper studies the variability, reliability and separability properties of FW-corrected DTI in the healthy human brain. We explore different methodologies used to correct the DTI for the FW compartment, depending on whether the diffusion-weighted MR acquisitions are single-or multiple-shell-based, and evaluate them using three publicly available databases acquired in an inter-session or inter-scanner manner. Our study has shown that one should not only look for the maximal repeatability (or reproducibility) of FW-corrected DTI measures in the brain studies, but also assess the reliability and separability indices, as it has been particularly observed with the FW-corrected MD parameter using the single-shell variational method. Importantly, how to correct the DTI for the FW is of great importance – the behaviour of the single-shell method appears to be data-dependent, with questionable enhancement in variability and reliability, as well as the FW-corrected MD parameter being anatomically non-meaningful. The multiple-shell FW-correction contraction-based technique has shown a reduced reproducibility and reliability of the measures compared to the standard DTI, the results being consistent across the evaluated data. As a conclusive remark, the FW correction to DTI should not be considered in terms of conceivable “improvement” in the repeatability (or reproducibility) and reliability, but rather as a methodology that provides information about the probed tissue, which the standard DTI partially hides. However, the choice of the FW-correction scheme should be made to avoid impacting the reproducibility, reliability, and accuracy of DTI-based parameters. As a solution, we suggest employing a customised FW-correction scheme, i.e., estimating the FWVF externally, correcting the diffusion-weighted MR signal for the FW, and then re-estimating the DTI using a standard procedure. Our experiments have provided evidence that this strategy improves the repeatability and reproducibility while not (significantly) affecting the reliability of the FW-corrected measures compared to the multiple-shell FW-correction of bi-tensor representation, while maintaining a relatively low error in the estimated MD and FA parameters.

## Acknowledgements

This work was funded by Junta de Castilla y León and Fondo Social Europeo Plus (FSE+) under research grant VA156P24. This work was supported by the Ministerio de Ciencia e Innovación of Spain with the research grant PID2021-124407NB-I00. Tomasz Pieciak acknowledges the Polish National Agency for Academic Exchange for grant PPN/BEK/2019/1/00421 under the Bekker programme and the Ministry of Science and Higher Education (Poland) under the scholarship for outstanding young scientists (692/STYP/13/2018). Guillem París was funded by the Consejería de Educación de Castilla y León and the European Social Fund through the “Ayudas para financiar la contratación predoctoral de personal investigador - Orden EDU/1100/201712/12” program.

The authors acknowledge Dr. Nico Lehmann for sharing the Magdeburg database.

## Ethics statement

Ethics approval were waived for this study due to the use of external data.

## Data availability

Data comes from the papers by Koller et al. (2021), Lehmann et al. (2021) and Tong et al. (2020).

## Declaration of generative AI

The authors did not use generative AI in writing of this manuscript.

## Footnotes

1 Percent point.

2 DiffKDE is a computational method used to estimate the probability density function and should not be confused with diffusion-weighted MR technique.

## References

Aja-Fernández, S., Vegas-Sánchez-Ferrero, G. (2016). Statistical analysis of noise in MRI. Springer International Publishing.

Albi, A., Pasternak, O., Minati, L., Marizzoni, M., Bartrés-Faz, D., Bargallo, N., et al. (2017). Free water elimination improves test–retest reproducibility of diffusion tensor imaging indices in the brain: A longitudinal multisite study of healthy elderly subjects. Human brain mapping, 38(1), 12–26.

Ades-Aron, B., Coelho, S., Lemberskiy, G., Veraart, J., Baete, S. H., Shepherd, T. M., et al. (2025). Denoising Improves Cross-Scanner and Cross-Protocol Test–Retest Reproducibility of Diffusion Tensor and Kurtosis Imaging. Human Brain Mapping, 46(4), e70142.

Andersson, J. L., Skare, S., Ashburner, J. (2003). How to correct susceptibility distortions in spin-echo echo-planar images: application to diffusion tensor imaging. Neuroimage, 20(2), 870–888.

Andersson, J. L., Graham, M. S., Zsoldos, E., Sotiropoulos, S. N. (2016). Incorporating outlier detection and replacement into a non-parametric framework for movement and distortion correction of diffusion MR images. Neuroimage, 141, 556–572.

Basser, P. J., Mattiello, J., LeBihan, D. (1994). Estimation of the effective self-diffusion tensor from the NMR spin echo. Journal of Magnetic Resonance, Series B, 103(3), 247–254.

Bergamino, M., Walsh, R. R., Stokes, A. M. (2021). Free-water diffusion tensor imaging improves the accuracy and sensitivity of white matter analysis in Alzheimer’s disease. Scientific reports, 11(1), 6990.

Bergmann, Ø., Henriques, R., Westin, C. F., Pasternak, O. (2020). Fast and accurate initialization of the free-water imaging model parameters from multi-shell diffusion MRI. NMR in Biomedicine, 33(3), e4219.

Boudreau, M., Karakuzu, A., Boré, A., Pinsard, B., Zelenkovski, K., Alonso-Ortiz, E., et al. (2025). Longitudinal reproducibility of brain and spinal cord quantitative MRI biomarkers. Imaging Neuroscience, 3, imag_a_00409.

Botev, Z. I., Grotowski, J. F., Kroese, D. P. (2010). Kernel density estimation via diffusion (2010). The Annals of Statistics, 38(5), 2916–2957.

Carreira Figueiredo, I., Borgan, F., Pasternak, O., Turkheimer, F. E., Howes, O. D. (2022). White-matter free-water diffusion MRI in schizophrenia: a systematic review and meta-analysis. Neuropsychopharmacology, 47(7), 1413–1420.

Chad, J. A., Pasternak, O., Salat, D. H., Chen, J. J. (2018). Re-examining age-related differences in white matter microstructure with free-water corrected diffusion tensor imaging. Neurobiology of aging, 71, 161–170.

Chad, J. A., Sochen, N., Chen, J. J., Pasternak, O. (2023). Implications of fitting a two-compartment model in single-shell diffusion MRI. Physics in Medicine & Biology, 68(21), 215012.

Chang, K., Burke, L., LaPiana, N., Howlett, B., Hunt, D., Dezelar, M., et al. (2024). Free water elimination tractometry for aging brains. BioRxiv. 10.1101/2024.11.10.622861

Correia, M. M., Henriques, R. N., Golub, M., Winzeck, S., Nunes, R. G. (2024). The trouble with free-water elimination using single-shell diffusion MRI data: A case study in ageing. Imaging Neuroscience, 2, 1–17.

Duan, F., Zhao, T., He, Y., Shu, N. (2015). Test–retest reliability of diffusion measures in cerebral white matter: A multiband diffusion MRI study. Journal of Magnetic Resonance Imaging, 42(4), 1106–1116.

Grech-Sollars, M., Hales, P. W., Miyazaki, K., Raschke, F., Rodriguez, D., Wilson, M., et al. (2015). Multicentre reproducibility of diffusion MRI parameters for clinical sequences in the brain. NMR in Biomedicine, 28(4), 468–485.

Guadilla, I., Fouto, A. R., Ruiz-Tagle, A., Esteves, I., Caetano, G., Silva, N. A., (2025). White matter alterations in episodic migraine without aura patients assessed with diffusion MRI: effect of free water correction. The Journal of Headache and Pain, 26(1), 1–13.

Golub, M., Henriques, R. N., Nunes, R. G. (2021). Free-water DTI estimates from single b-value data might seem plausible but must be interpreted with care. Magnetic resonance in medicine, 85(5), 2537–2551.

Hastie, T., Tibshirani, R., Friedman, J. H., Friedman, J. H. (2009). The elements of statistical learning: data mining, inference, and prediction, Springer

Henriques, N. R., Rokem, A., Garyfallidis, E., St-Jean, S., Peterson, E.T. and Correia, M.M. (2017). [Re] Optimization of a free water elimination two-compartment model for diffusion tensor imaging, ReScience, 3(1), #2.

Hoy, A. R., Koay, C. G., Kecskemeti, S. R., Alexander, A. L. (2014). Optimization of a free water elimination two-compartment model for diffusion tensor imaging. Neuroimage, 103, 323–333.

Jakab, A., Tuura, R., Kellenberger, C., Scheer, I. (2017). In utero diffusion tensor imaging of the fetal brain: a reproducibility study. NeuroImage: Clinical, 15, 601–612.

Kellner, E., Dhital, B., Kiselev, V. G., Reisert, M. (2016). Gibbs-ringing artifact removal based on local subvoxel-shifts. Magnetic resonance in medicine, 76(5), 1574–1581.

Koay, C. G., Chang, L. C., Carew, J. D., Pierpaoli, C., Basser, P. J. (2006). A unifying theoretical and algorithmic framework for least squares methods of estimation in diffusion tensor imaging. Journal of magnetic resonance, 182(1), 115–125.

Koller, K., Rudrapatna, U., Chamberland, M., Raven, E. P., Parker, G. D., Tax, C. M., et al. (2021). MICRA: Microstructural image compilation with repeated acquisitions. Neuroimage, 225, 117406.

Laguna, P. A. L., Combes, A. J., Streffer, J., Einstein, S., Timmers, M., Williams, S. C., Dell’Acqua, F. (2020). Reproducibility, reliability and variability of FA and MD in the older healthy population: A test-retest multiparametric analysis. NeuroImage: Clinical, 26, 102168.

Le Bihan, D., Johansen-Berg, H. (2012). Diffusion MRI at 25: exploring brain tissue structure and function. Neuroimage, 61(2), 324–341.

Lehmann, N., Aye, N., Kaufmann, J., Heinze, H. J., Düzel, E., Ziegler, G., Taubert, M. (2021). Longitudinal reproducibility of neurite orientation dispersion and density imaging (NODDI) derived metrics in the white matter. Neuroscience, 457, 165–185.

Liu, Q., Ning, L., Shaik, I. A., Liao, C., Gagoski, B., Bilgic, B., et al. (2024). Reduced cross-scanner variability using vendor-agnostic sequences for single-shell diffusion MRI. Magnetic Resonance in Medicine, 92(1), 246–256.

Lyall, A. E., Pasternak, O., Robinson, D. G., Newell, D., Trampush, J. W., Gallego, J. A., et al. (2018). Greater extracellular free-water in first-episode psychosis predicts better neurocognitive functioning. Molecular psychiatry, 23(3), 701–707.

Maillard, P., Fletcher, E., Singh, B., Martinez, O., Johnson, D. K., Olichney, J. M., et al. (2019). Cerebral white matter free water: A sensitive biomarker of cognition and function. Neurology, 92(19), e2221–e2231

Melzer, T. R., Keenan, R. J., Leeper, G. J., Kingston-Smith, S., Felton, S. A., Green, S. K., et al. (2020). Test-retest reliability and sample size estimates after MRI scanner relocation. Neuroimage, 211, 116608.

Metzler-Baddeley, C., O’Sullivan, M. J., Bells, S., Pasternak, O., Jones, D. K. (2012). How and how not to correct for CSF-contamination in diffusion MRI. Neuroimage, 59(2), 1394–1403.

Merisaari, H., Tuulari, J. J., Karlsson, L., Scheinin, N. M., Parkkola, R., Saunavaara, J., et al. (2019). Test-retest reliability of diffusion tensor imaging metrics in neonates. NeuroImage, 197, 598–607.

Molina-Romero, M., Wiestler, B., Gómez, P. A., Menzel, M. I., Menze, B. H. (2018). Deep learning with synthetic diffusion MRI data for free-water elimination in glioblastoma cases. In International Conference on Medical Image Computing and Computer-Assisted Intervention (pp. 98–106). Cham: Springer International Publishing.

Mori, S., Wakana, S., Van Zijl, P. C., Nagae-Poetscher, L. M. (2005). MRI atlas of human white matter. Elsevier.

Nakaya, M., Sato, N., Matsuda, H., Maikusa, N., Shigemoto, Y., Sone, D., et al. (2022). Free water derived by multi-shell diffusion MRI reflects tau/neuroinflammatory pathology in Alzheimer’s disease. Alzheimer’s & Dementia: Translational Research & Clinical Interventions, 8(1), e12356.

Nemmi, F., Levardon, M., Péran, P. (2022). Brain-age estimation accuracy is significantly increased using multishell free-water reconstruction. Human Brain Mapping, 43(7), 2365–2376.

Ofori, E., Pasternak, O., Planetta, P. J., Li, H., Burciu, R. G., Snyder, A. F., et al. (2015). Longitudinal changes in free-water within the substantia nigra of Parkinson’s disease. Brain, 138(8), 2322–2331.

Parker, D., Ould Ismail, A. A., Wolf, R., Brem, S., Alexander, S., Hodges, W., et al. (2020). Freewater estimatoR using iNtErpolated iniTialization (FERNET): Characterizing peritumoral edema using clinically feasible diffusion MRI data. Plos one, 15(5), e0233645.

Pasternak, O., Sochen, N., Gur, Y., Intrator, N., Assaf, Y. (2009). Free water elimination and mapping from diffusion MRI. Magnetic resonance in medicine. 62(3), 717–730.

Pasternak, O., Shenton, M. E., Westin, C. F. (2012). Estimation of extracellular volume from regularized multi-shell diffusion MRI. In International conference on medical image computing and computer-assisted intervention (pp. 305–312). Berlin, Heidelberg: Springer Berlin Heidelberg.

Pieciak, T., Rabanillo-Viloria, I., Aja-Fernández, S. (2018). Bias correction for non-stationary noise filtering in MRI. In 2018 IEEE 15th International Symposium on Biomedical Imaging (ISBI 2018) (pp. 307–310). IEEE.

Pieciak, T., Aja-Fernandez, S., Vegas-Sanchez-Ferrero, G. (2017). Non-stationary Rician noise estimation in parallel mri using a single image: a variance-stabilizing approach. IEEE transactions on pattern analysis and machine intelligence, 39(10), 2015–2029.

Pieciak, T., París, G., Beck, D., Maximov, I. I., Tristán-Vega, A., de Luis-García, R., Westlye, L.T., Aja-Fernández, S. (2023). Spherical means-based free-water volume fraction from diffusion MRI increases non-linearly with age in the white matter of the healthy human brain. Neuroimage, 279, 120324.

Pierpaoli, C., Jones, D. K. (2004). Removing CSF contamination in brain DT-MRIs by using a two-compartment tensor model. In International Society for Magnetic Resonance in Medicine Meeting, 1215.

Rydhög, A. S., Szczepankiewicz, F., Wirestam, R., Ahlgren, A., Westin, C. F., Knutsson, L., Pasternak, O. (2017). Separating blood and water: Perfusion and free water elimination from diffusion MRI in the human brain. Neuroimage, 156, 423–434.

Shahim, P., Holleran, L., Kim, J. H., Brody, D. L. (2017). Test-retest reliability of high spatial resolution diffusion tensor and diffusion kurtosis imaging. Scientific Reports, 7(1), 11141.

Sheather, S. J., Jones, M. C. (1991). A reliable data-based bandwidth selection method for kernel density estimation. Journal of the Royal Statistical Society: Series B (Methodological), 53(3), 683–690.

Silverman, B. W. (2018). Density estimation for statistics and data analysis. Routledge.

Smith, S. M., Jenkinson, M., Woolrich, M. W., Beckmann, C. F., Behrens, T. E., Johansen-Berg, H., et al. (2004). Advances in functional and structural MR image analysis and implementation as FSL. Neuroimage, 23, S208–S219.

Tong, Q., He, H., Gong, T., Li, C., Liang, P., Qian, T., et al. (2020). Multicenter dataset of multi-shell diffusion MRI in healthy traveling adults with identical settings. Scientific Data, 7(1), 157.

Tristán-Vega, A., París, G., de Luis-García, R., Aja-Fernández, S. (2022). Accurate free-water estimation in white matter from fast diffusion MRI acquisitions using the spherical means technique. Magnetic Resonance in Medicine, 87(2), 1028–1035.

Tustison, N. J., Avants, B. B., Cook, P. A., Zheng, Y., Egan, A., Yushkevich, P. A., Gee, J. C. (2010). N4ITK: improved N3 bias correction. IEEE transactions on medical imaging, 29(6), 1310–1320.

Van Essen, D. C., Smith, S. M., Barch, D. M., Behrens, T. E., Yacoub, E., Ugurbil, K., Wu-Minn HCP Consortium. (2013). The WU-Minn human connectome project: an overview. Neuroimage, 80, 62–79.

Venkatraman, V. K., Gonzalez, C. E., Landman, B., Goh, J., Reiter, D. A., An, Y., Resnick, S. M. (2015). Region of interest correction factors improve reliability of diffusion imaging measures within and across scanners and field strengths. Neuroimage, 119, 406–416.

Veraart, J., Novikov, D. S., Christiaens, D., Ades-Aron, B., Sijbers, J., Fieremans, E. (2016a). Denoising of diffusion MRI using random matrix theory. Neuroimage, 142, 394–406.

Veraart, J., Fieremans, E., Novikov, D. S. (2016b). Diffusion MRI noise mapping using random matrix theory. Magnetic resonance in medicine, 76(5), 1582–1593.

Westin, C. F., Maier, S. E., Mamata, H., Nabavi, A., Jolesz, F. A., Kikinis, R. (2002). Processing and visualization for diffusion tensor MRI. Medical image analysis, 6(2), 93–108.

Weninger, L., Koppers, S., Na, C. H., Juetten, K., Merhof, D. (2020). Free-water correction in diffusion mri: a reliable and robust learning approach. In Computational Diffusion MRI: MICCAI Workshop, Shenzhen, China, October 2019 (pp. 91–99). Springer International Publishing.

Zhong, J., Liu, X., Hu, Y., Xing, Y., Ding, D., Ge, X., et al. (2024). Robustness of Quantitative Diffusion Metrics from Four Models: A Prospective Study on the Influence of Scan-Rescans, Voxel Size, Coils, and Observers. Journal of Magnetic Resonance Imaging, 60(4), 1470–1483.

Zuo, X. N., Xu, T., Milham, M. P. (2019). Harnessing reliability for neuroscience research. Nature human behaviour, 3(8), 768–771.

