## Supplementary materials for "Reproducibility and reliability of Free-Water-corrected Diffusion Tensor Imaging of the brain: Revisited"

### 1. Materials and methods

#### 1.1. Additional datasets

The acquisition setups for two additional databases were as follows:

- Magdeburg: The database includes longitudinal test-retest acquisitions from a single scanner separated by four weeks (Lehmann et al., 2021). Twenty-nine healthy volunteers (24F/5M, aged 19–35/19–30) were scanned using a 3T MAGNETOM Prisma scanner (Siemens Healthcare, Erlangen, Germany) equipped with a 64-channel head coil. Acquisition protocol: single-shot spin-echo echo planar imaging sequence, parallel acquisition using the Generalized Autocalibrating Partially Parallel Acquisition (GRAPPA) at factor 2, multi-band acquisition at multi-band factor 2, AP phase-encoding direction, TR: 4970 ms, TE: 74 ms, FOV: 208 × 208 $mm^2$, matrix size: 130 × 130, voxel size: 1.6 × 1.6 × 1.6 $mm^3$, *b*-values: (1000, 2000, 3000) $s/mm^2$ with (38, 76, 114) gradient directions, respectively, 14 non-diffusion-weighted scans in AP direction and 9 non-diffusion-weighted scans in PA direction.

- ZJU: The database includes inter-scanner volumes from 10 centres (Tong et al., 2020). Three healthy volunteers (2F/1M, aged 23, 26 and 23) were scanned once in nine centres and three times at tenth centre. Each centre has been equipped with the same 3T MR MAGNETOM Prisma scanner (Siemens Healthcare, Erlangen, Germany) with maximal gradient strength of 80 mT/m. Acquisition protocol: simultaneous multi-slice (SMS) spin-echo echo planar imaging sequence, SMS factor 2, parallel GRAPPA at factor 2, AP phase-encoding direction, TR: 5400 ms, TE: 71 ms, $\Delta/\delta$: 34.4/15.9 ms, FOV: 220 × 220 $mm^2$, voxel size: 1.5 × 1.5 × 1.5 $mm^3$, *b*-values: (1000, 2000, 3000) $s/mm^2$ with 30 gradient directions per shell, and 6 non-diffusion-weighted scans.

- Multi *b*-value acquisition: A single healthy volunteer was scanned using a Siemens Trio 3T scanner (Siemens Healthcare, Erlangen, Germany) equipped with a 32-channel head coil (Hansen & Jespersen, 2016). Acquisition protocol: TR: 7200 ms, TE: 116 ms, TI: 2100 ms, $\Delta/\delta$: 58/29 ms, *b*-values ranges from 0 to 3000 $s/mm^2$ with the step size of 200 $s/mm^2$ and 33 gradient directions per shell, and a single non-diffusion-weighted MR scan.

#### 1.2. Data preprocessing

- Magdeburg: The database has been shared partially preprocessed, including susceptibility-induced, eddy current and head movements distortions with the FSL `topup` (Andersson et al., 2003; Smith et al., 2004) and FSL `eddy` (Andersson et al., 2016) tools. We additionally employed the MICRA preprocessing chain as follows: 1) noise removal *via* the MP-PCA (Veraart et al., 2016a; Veraart et al., 2016b), 2) Gibbs ringing artefacts correction (Kellner et al., 2016), 3) Rician bias correction (Pieciak et al., 2018), 4) B1 field inhomogeneity correction using the N4 algorithm (Tustison et al., 2010).

- ZJU: The database has been publicly shared in a preprocessed variant. The preprocessing pipeline covered: 1) noise removal using the MP-PCA (Veraart et al., 2016a; Veraart et al., 2016b), 2) Gibbs ringing artefacts correction (Kellner et al., 2016), 3) susceptibility-induced distortions estimation using the FSL `topup` (Andersson et al., 2003) and 4) head movements and eddy current distortions correction with the FSL `eddy` (Andersson et al., 2016). No further preprocessing was carried on.

- Multi-*b*-value acquisition: The data has been corrected for 1) noise using the MP-PCA approach (Veraart et al., 2016a; Veraart et al., 2016b) and 2) Rician bias (Pieciak et al., 2018).

### 2. Experimental results

The experimental results in this supplementary materials part include the following parameters computed for *in silico* data generated according to Eq. (1) in the main document and *in vivo* Magdeburg and ZJU databases:

1. FWVF parameters were estimated from single-shell data using the Bi-tensor-S method (Pasternak et al., 2009), and multiple-shell data with the SM (Tristán-Vega et al., 2022) and Bi-tensor-M (Hoy et al., 2014),

2. DTI measures were estimated from single- and multiple-shell data (Koay et al., 2006),

3. FW-corrected DTI measures were estimated from 1) single-shell data with Bi-tensor-S approach (Pasternak et al., 2009), 2) multiple-shell data with Bi-tensor-M (Hoy et al., 2014) and 3) the FW-DTI customized approach that computes DTI from FW-corrected diffusion-weighted MR signal, as defined by Eq. (5) in the main document. The FW-DTI covers both single- and multiple-shell data.

Additionally, the *in vivo* experiments cover computing the FWVF using the SM and Bi-tensor-M for multi-*b*-value acquisition data (Hansen & Jespersen, 2016).

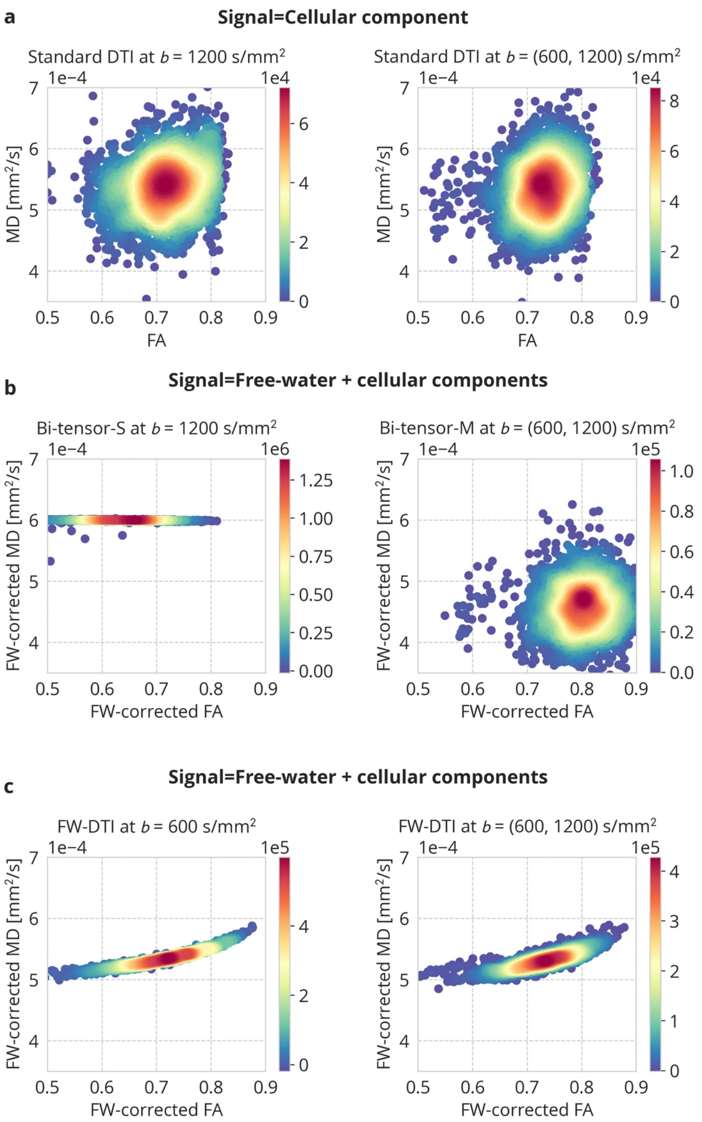


**Figure S1.** The 2D density plots illustrate the experimental results from an _*in silico*_ data under a single-fibre bundle configuration: **a)** the MD and FA parameters estimated using the standard DTI from noiseless reference data covering the cellular component only. The FW-corrected MD and FA estimated from the signal covering FW ($f=0.2$) and cellular components: **b)** Bi-tensor-S and Bi-tensor-M approaches, and **c)** FW-DTI customised scheme according to Eq. (5). In total, $15^3$ samples have been used to obtain a single 2D density plot.

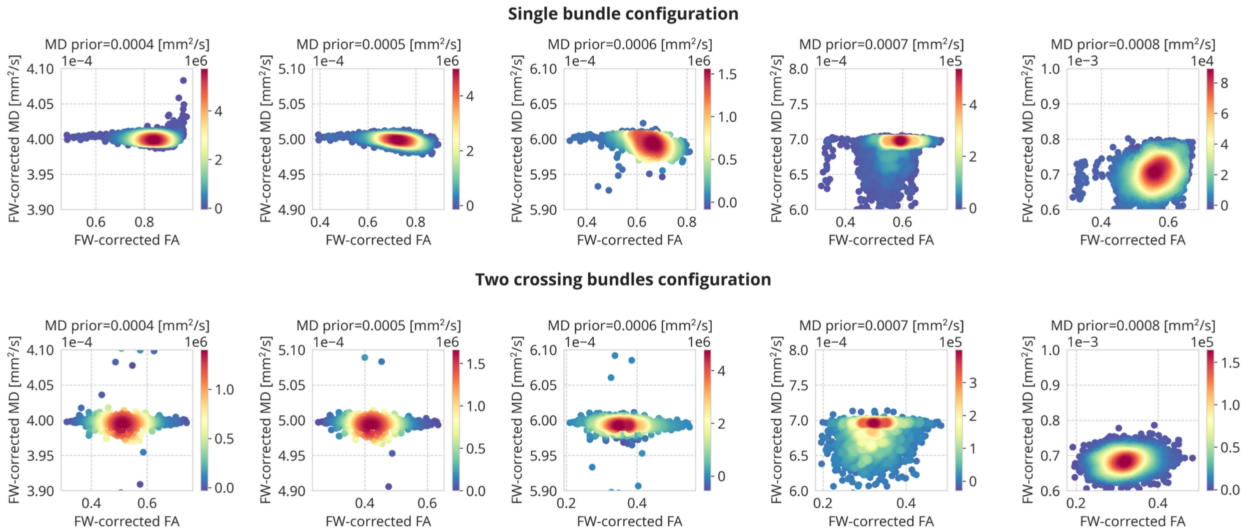


**Figure S2.** The 2D density plots illustrate estimated FW-corrected MD and FA parameters using the method by Pasternak et al. (2009) from _*in silico*_ _single-shell data_ generated at $b = 1200$ s/mm$^2$. Each figure covers the results under different tissue's MD prior. The top row presents the results for a single bundle configuration (i.e. $\alpha_2=0$ in Eq. (1)), while the bottom row demonstrates the results for two crossing bundles (i.e., $\alpha_1=\alpha_2=0.5$ in Eq. (1)). Both variants assume the FWVF $f=0.2$. In total, $15^3$ samples have been used to obtain a single 2D density plot.

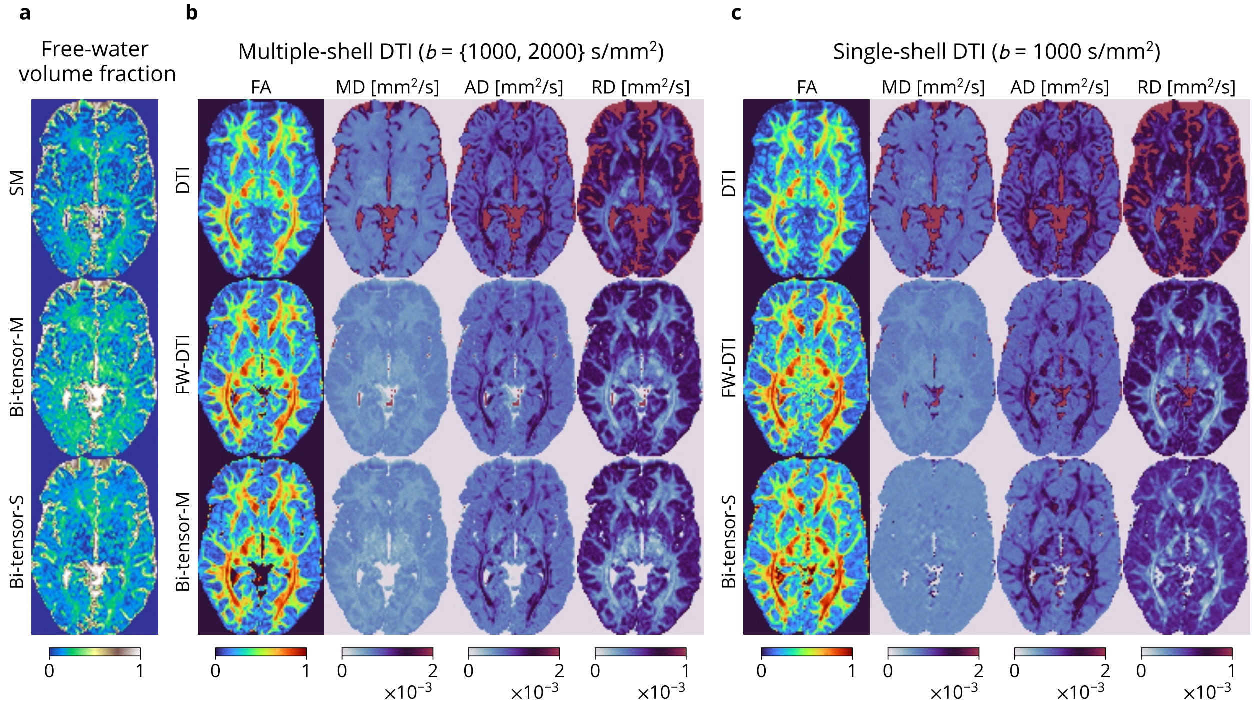


**Figure S3.** Estimated microstructural measures for a selected Magdeburg acquisition (av38, ses-1, slice: 40): **a)** FWVF estimated from multiple-shell data (SM and Bi-tensor-M approaches) and single-shell data (Bi-tensor-S), **b)** DTI-based measures estimated from multiple-shell data using a standard DTI, and two free water-correction methodologies: FW-DTI according to Eq. (5) and Bi-tensor-M, and **c)** DTI-based measures estimated from single-shell data with the standard DTI, FW-DTI and Bi-tensor-S. The FWVF parameter for FW-DTI was pre-estimated using the SM approach from multiple-shell data in both scenarios presented in panels b) and c).

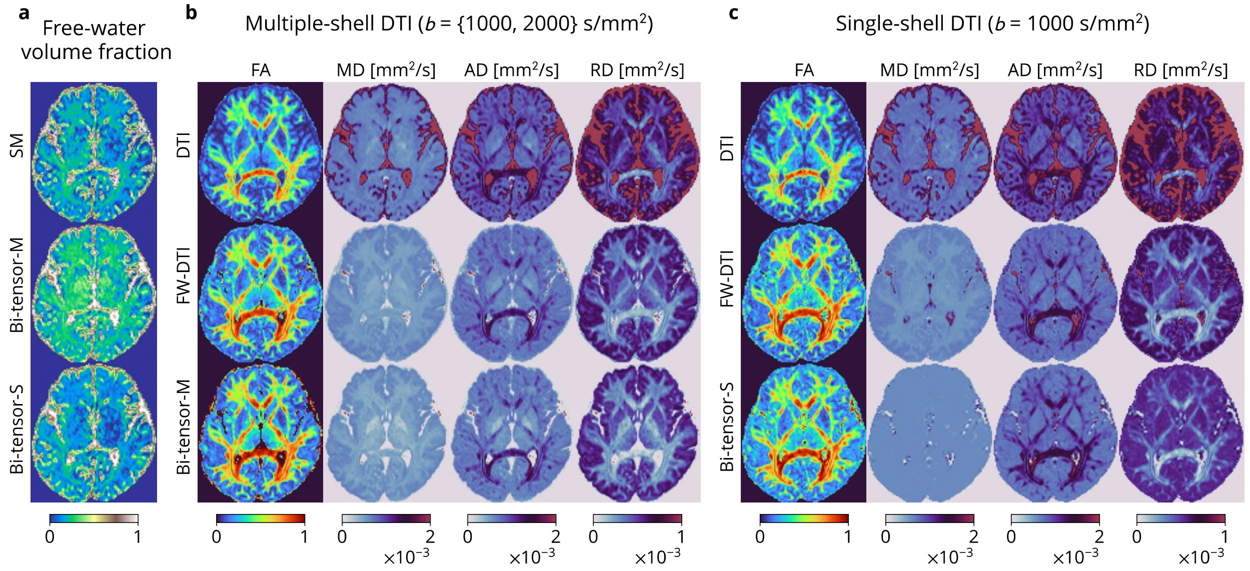


**Figure S4.** Estimated microstructural measures for a selected ZJU acquisition (sub-1, ses-c01r1, slice: 46).

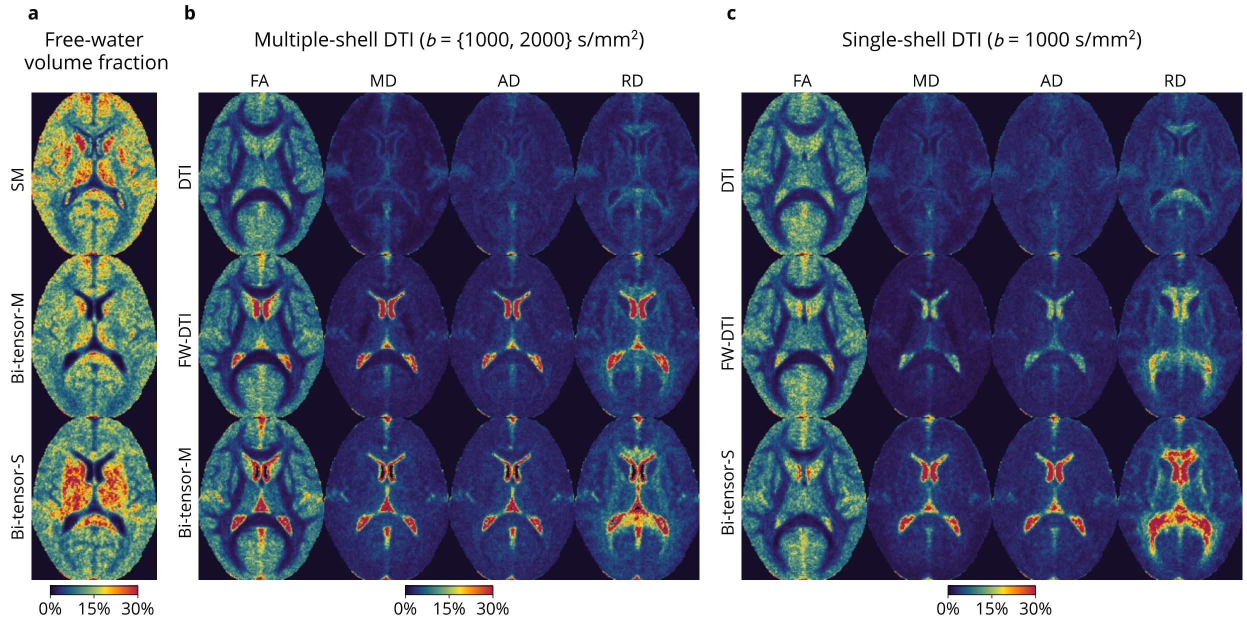


**Figure S5.** Inter-session longitudinal variability maps of the measures defined in the standard space for Magdeburg database according to the coefficient of variation defined by Eq. (8): **a)** FWVF estimated from multiple-shell data (SM and Bi-tensor-M approaches) and single-shell data (Bi-tensor-S), **b)** DTI-based measures estimated from multiple-shell data using a standard DTI, and two FW-correction methodologies: FW-DTI according to Eq. (5) and Bi-tensor-M, and **c)** DTI-based measures estimated from single-shell data with the standard DTI, FW-DTI and Bi-tensor-S. The FWVF parameter for FW-DTI was pre-estimated using the SM approach from multiple-shell data in both scenarios presented in panels b) and c).

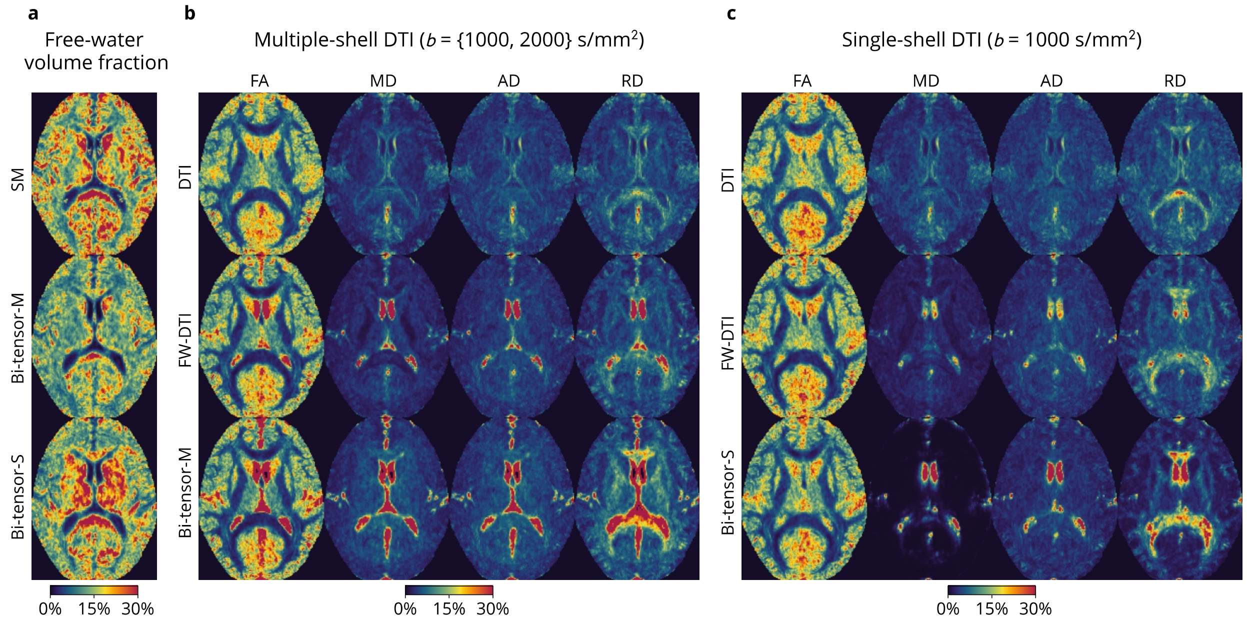


**Figure S6.** Inter-scanner variability maps of the measures defined in the standard space for ZJU database according to the coefficient of variation by Eq. (8).

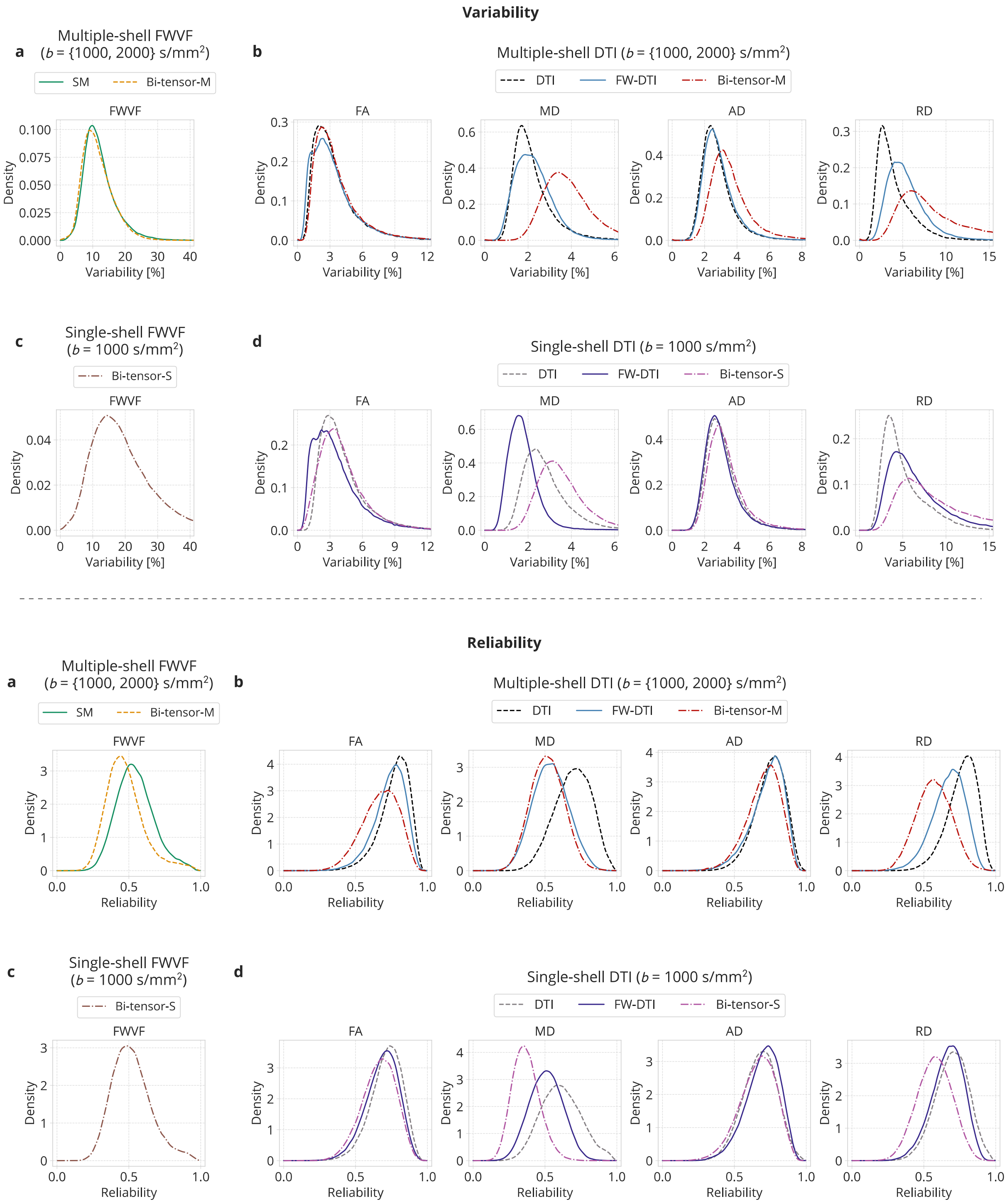


**Figure S7.** Inter-session longitudinal kernel density-based variability (top) and reliability (bottom) indices computed for Magdeburg database over the white matter area: **a)** FWVF estimated from multiple-shell data (SM and Bi-tensor-M approaches), **b)** DTI-based measures estimated from multiple-shell data using standard DTI, FW-DTI according to Eq. (5) and Bi-tensor-M, **c)** FWVF estimated from single-shell data (Bi-tensor-S) and **d)** DTI-based measures estimated from single-shell data with the standard DTI, FW-DTI and Bi-tensor-S. The FWVF parameter for FW-DTI was pre-estimated from multiple-shell data using the SM approach in both scenarios presented in panels b) and d).

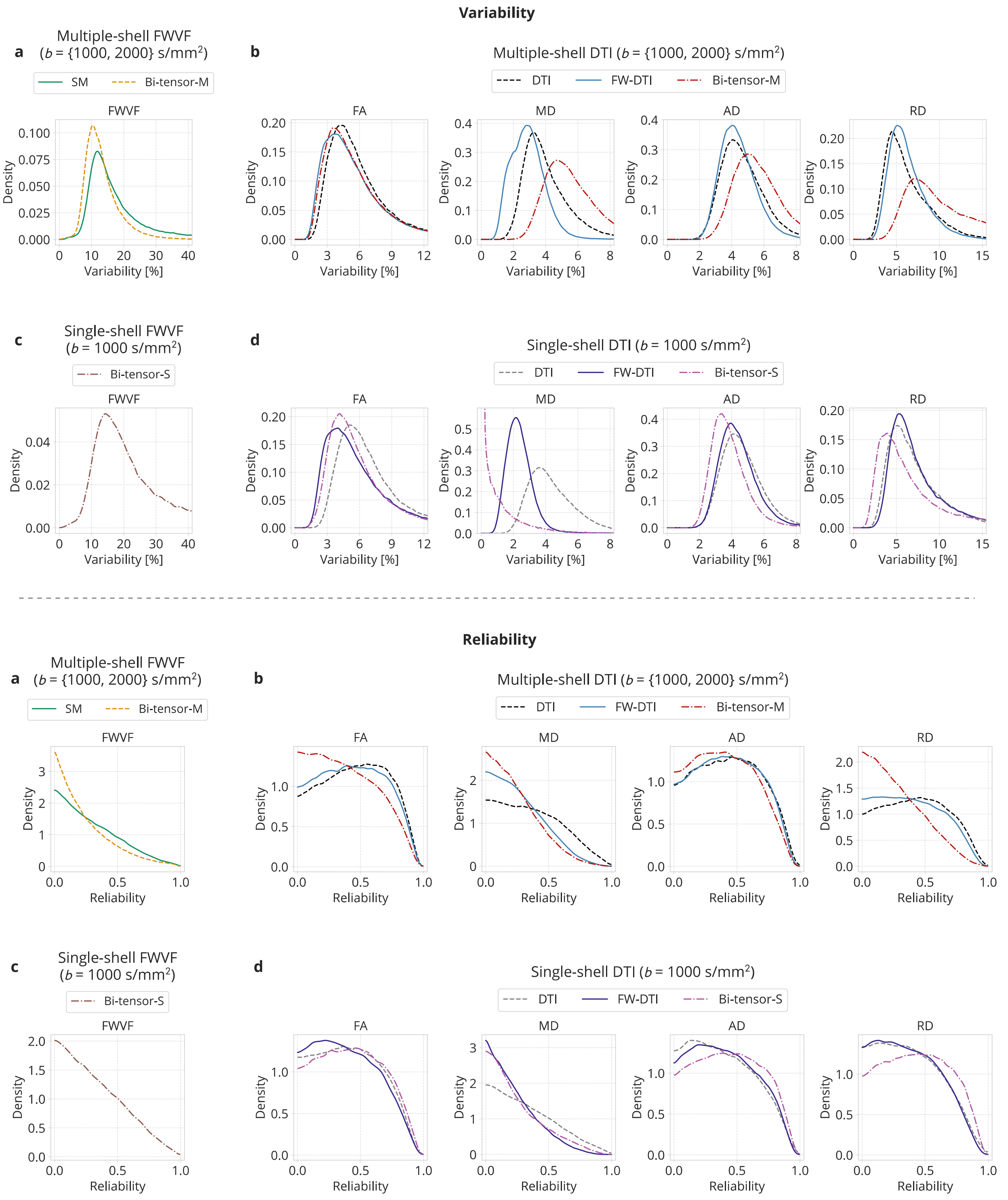


**Figure S8.** Inter-scanner kernel density-based variability (top) and reliability (bottom) indices computed for <u>ZJU database</u> over the white matter area.

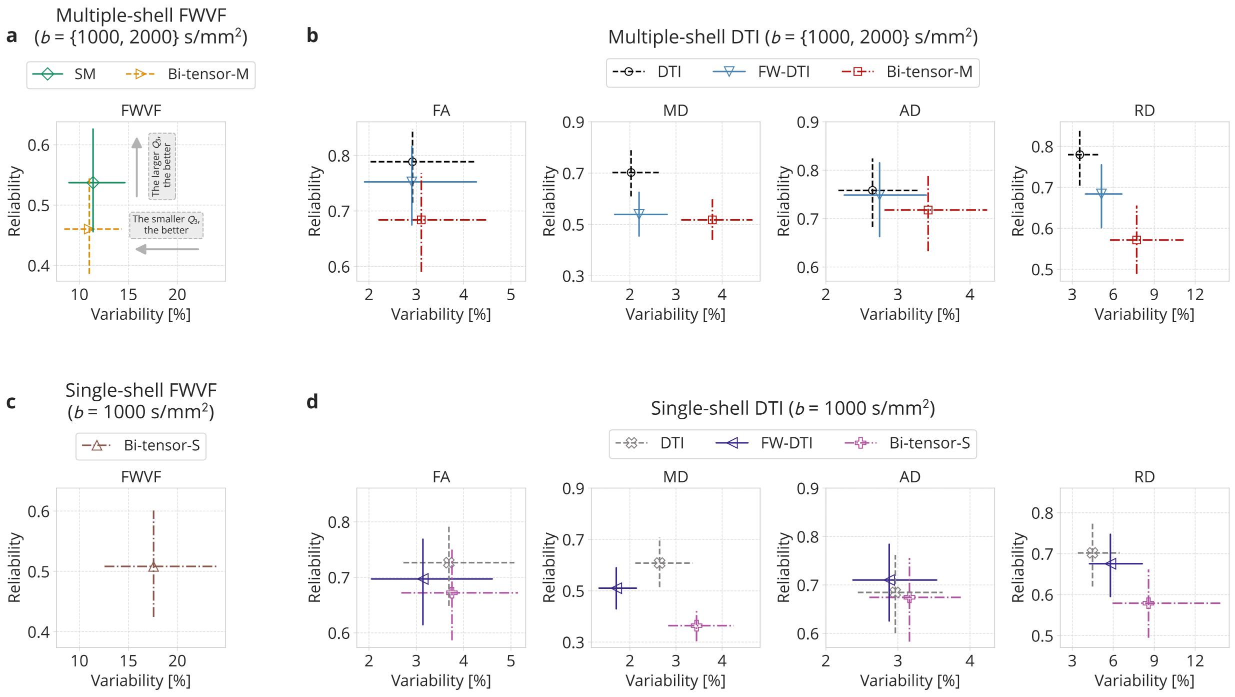


**Figure S9.** Inter-session longitudinal reliability versus variability plots computed for Magdeburg database over the white matter area: **a)** FWVF estimated from multiple-shell data (SM and Bi-tensor-M approaches), **b)** DTI-based measures estimated from multiple-shell data using standard DTI, FW-DTI according to Eq. (5) and Bi-tensor-M, **c)** FWVF estimated from single-shell data (Bi-tensor-S) and **d)** DTI-based measures estimated from single-shell data with the standard DTI, FW-DTI and Bi-tensor-S. The FWVF parameter for FW-DTI was pre-estimated from multiple-shell data using the SM approach in both scenarios presented in panels b) and d). The markers present median values calculated over the white matter area, while the horizontal and vertical lines represent distances between the first $Q_1 = 0.25$ (25th percentile) and third $Q_3 = 0.75$ (75th percentile) quartiles. The horizontal and vertical ranges for the plots representing multiple- and single-shell equivalent cases have been fixed.

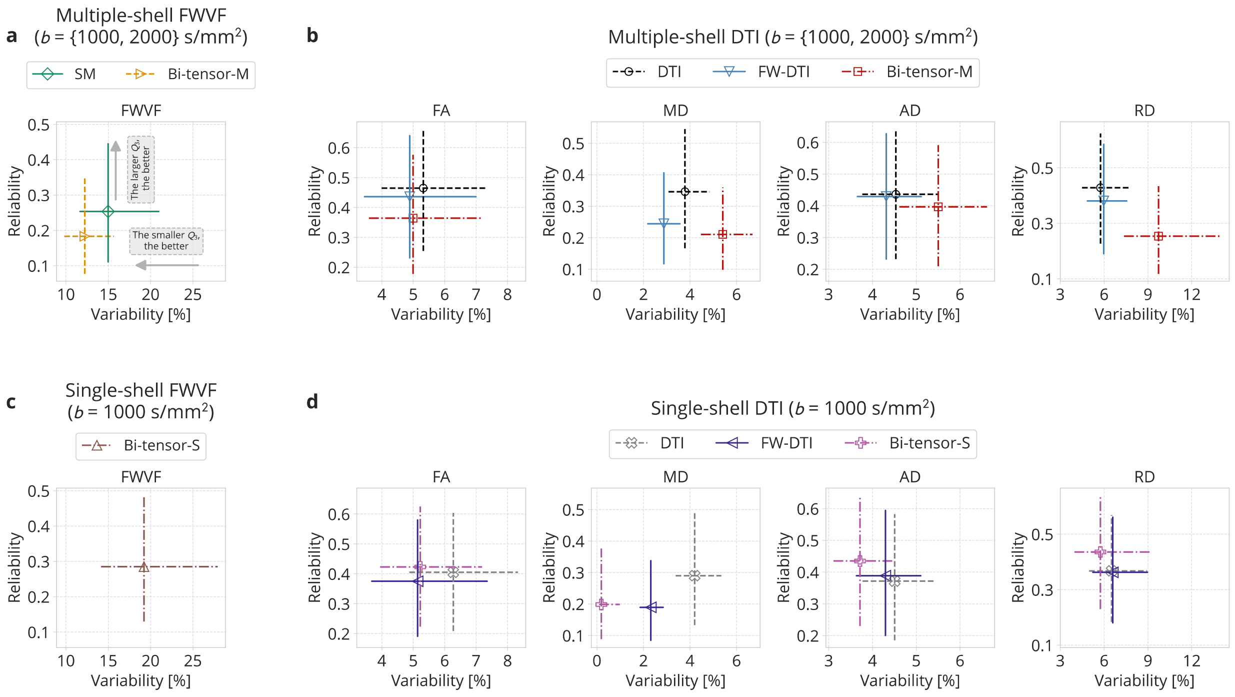


**Figure S10.** Inter-scanner reliability versus variability plots computed for ZJU database over the white matter area.

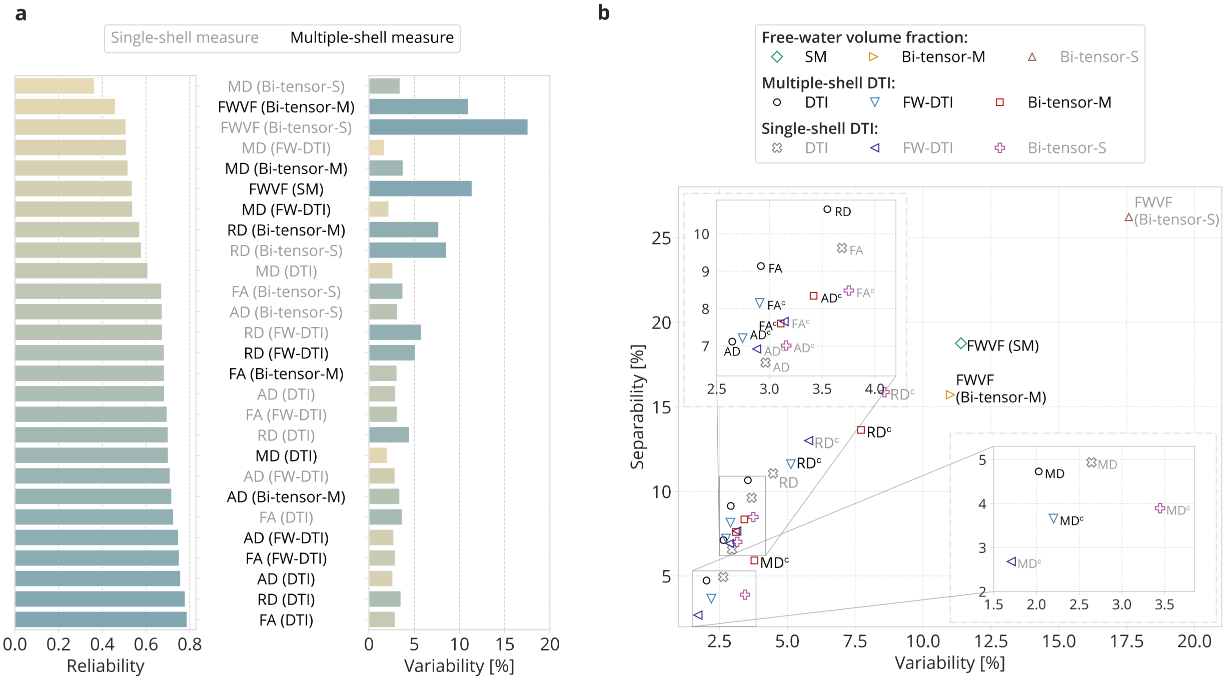


**Figure S11. a)** Box-plots presenting inter-session longitudinal median reliability and median variability indices computed from the Magdeburg database over the white matter area. The colours in the variability box-plot reflect the ordering according to the reliability indice value. **b)** A diagram representing the separability indice as a function of the variability indice. A single marker refers to the median variability and median separability values calculated over the white matter area in the standard space. Multiple-shell measures are annotated with black text, while the single-shell measures are in grey. The FW-corrected DTI measures are annotated with a subscript "c".

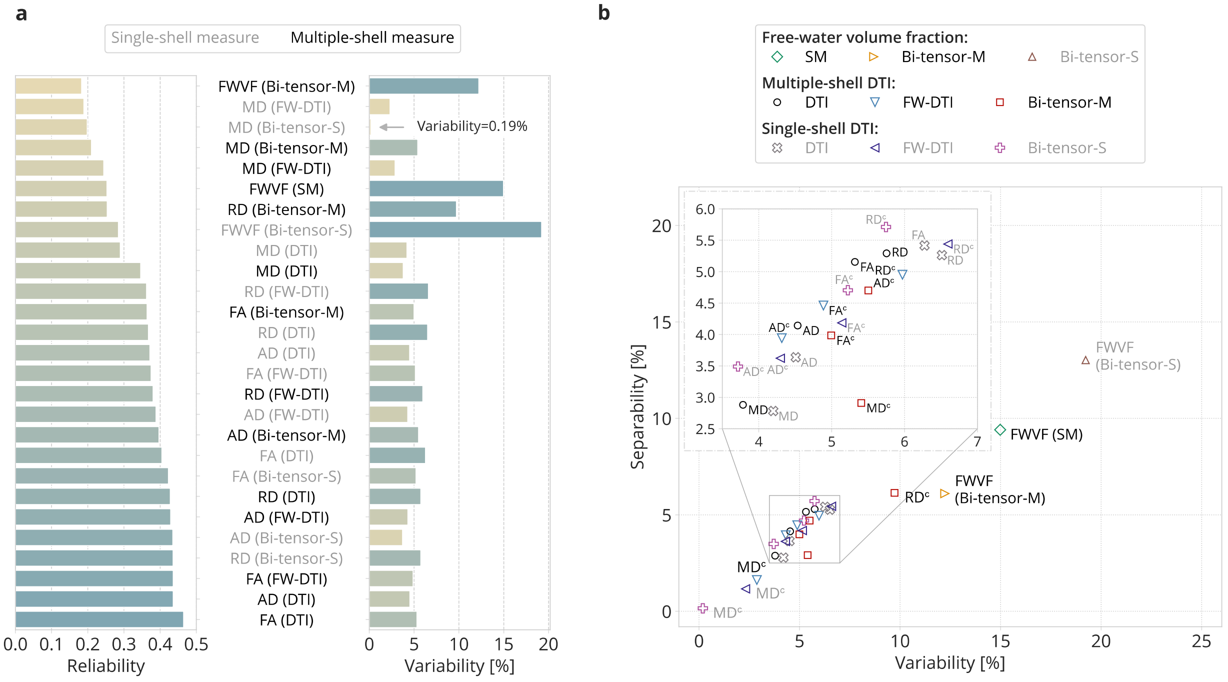


**Figure S12.** Box-plots presenting inter-scanner median reliability and median variability indexes computed from the ZJU database.

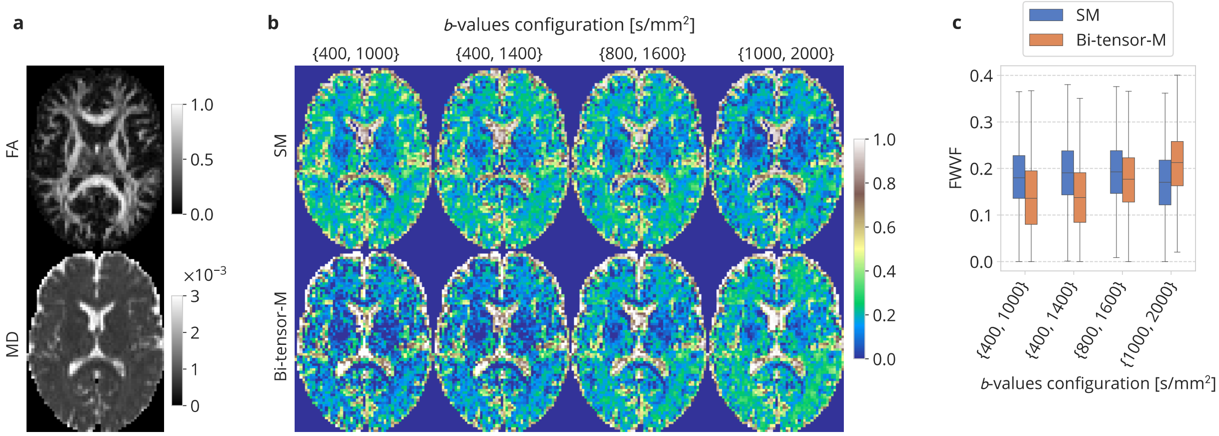


**Figure S13. a)** The standard DTI-based FA and MD estimated from multi-*b*-value data at $b = 1000$ s/mm$^2$ (Hansen and Jespersen, 2016). Changes in the FWVF parameter estimated from multi-*b*-value data using the SM and Bi-tensor-M under different *b*-values configurations: **b)** visual inspection of the measures and **c)** box-plots representing the quantified FWVF values over the white matter area. The white matter was determined by thresholding the standard DTI-based FA parameter over 0.3.

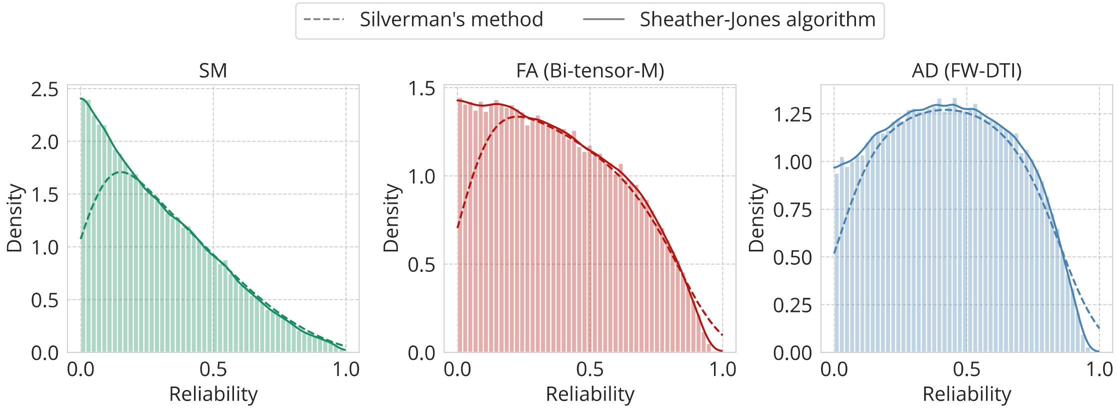


**Figure S14.** Comparison between kernel density-based plots computed using a standard KDE with Silverman's bandwidth selection method (Silverman, 1986) and DiffKDE with the improved Sheather-Jones algorithm used for bandwidth selection (Sheather and Jones, 1991; Botev et al., 2010). The plots demonstrate the inter-scanner reliability of the measures computed for the ZJU database and have been selected to exemplify different shapes of the density plots appeared in our experiments.

### References


Andersson, J. L., Skare, S., Ashburner, J. (2003). How to correct susceptibility distortions in spin-echo echo-planar images: application to diffusion tensor imaging. Neuroimage, 20(2), 870-888.

Andersson, J. L., Graham, M. S., Zsoldos, E., Sotiropoulos, S. N. (2016). Incorporating outlier detection and replacement into a non-parametric framework for movement and distortion correction of diffusion MR images. Neuroimage, 141, 556-572.

Botev, Z. I., Grotowski, J. F., Kroese, D. P. (2010). Kernel density estimation via diffusion. The Annals of Statistics, 38(5), 2916-2957.

Hansen, B., Jespersen, S. N. (2016). Data for evaluation of fast kurtosis strategies, b-value optimization and exploration of diffusion MRI contrast. Scientific data, 3(1), 1-5.

Hoy, A. R., Koay, C. G., Kecskemeti, S. R., Alexander, A. L. (2014). Optimization of a free water elimination two-compartment model for diffusion tensor imaging. Neuroimage, 103, 323-333.

Kellner, E., Dhital, B., Kiselev, V. G., Reisert, M. (2016). Gibbs-ringing artifact removal based on local subvoxel-shifts. Magnetic resonance in medicine, 76(5), 1574-1581.

Koay, C. G., Chang, L. C., Carew, J. D., Pierpaoli, C., Basser, P. J. (2006). A unifying theoretical and algorithmic framework for least squares methods of estimation in diffusion tensor imaging. Journal of magnetic resonance, 182(1), 115-125.

Lehmann, N., Aye, N., Kaufmann, J., Heinze, H. J., Düzel, E., Ziegler, G., Taubert, M. (2021). Longitudinal reproducibility of neurite orientation dispersion and density imaging (NODDI) derived metrics in the white matter. Neuroscience, 457, 165-185.

Pasternak, O., Sochen, N., Gur, Y., Intrator, N., Assaf, Y. (2009). Free water elimination and mapping from diffusion MRI. Magnetic resonance in medicine. 62(3), 717-730.

Pieciak, T., Rabanillo-Viloria, I., Aja-Fernández, S. (2018). Bias correction for non-stationary noise filtering in MRI. In 2018 IEEE 15th International Symposium on Biomedical Imaging (ISBI 2018) (pp. 307-310). IEEE.

Sheather, S.J., Jones, M. C. (1991). A reliable data-based bandwidth selection method for kernel density estimation. Journal of the Royal Statistical Society: Series B (Methodological), 53(3), 683-690.

Silverman, B.W. (1986). Density Estimation for Statistics and Data Analysis. London: Chapman & Hall/CRC.

Smith, S. M., Jenkinson, M., Woolrich, M. W., Beckmann, C. F., Behrens, T. E., Johansen-Berg, H., et al. (2004). Advances in functional and structural MR image analysis and implementation as FSL. Neuroimage, 23, S208-S219.

Tong, Q., He, H., Gong, T., Li, C., Liang, P., Qian, T., et al. (2020). Multicenter dataset of multi-shell diffusion MRI in healthy traveling adults with identical settings. Scientific Data, 7(1), 157.

Tristán-Vega, A., París, G., de Luis-García, R., Aja-Fernández, S. (2022). Accurate free-water estimation in white matter from fast diffusion MRI acquisitions using the spherical means technique. Magnetic Resonance in Medicine, 87(2), 1028-1035.

Tustison, N. J., Avants, B. B., Cook, P. A., Zheng, Y., Egan, A., Yushkevich, P. A., Gee, J. C. (2010). N4ITK: improved N3 bias correction. IEEE transactions on medical imaging, 29(6), 1310-1320.

Veraart, J., Novikov, D. S., Christiaens, D., Ades-Aron, B., Sijbers, J., Fieremans, E. (2016a). Denoising of diffusion MRI using random matrix theory. Neuroimage, 142, 394-406.

Veraart, J., Fieremans, E., Novikov, D. S. (2016b). Diffusion MRI noise mapping using random matrix theory. Magnetic resonance in medicine, 76(5), 1582-1593.
